# Elementary vectors and autocatalytic sets for computational models of cellular growth

**DOI:** 10.1101/2021.10.31.466640

**Authors:** Stefan Müller, Diana Széliová, Jürgen Zanghellini

## Abstract

Traditional (genome-scale) metabolic models of cellular growth involve an approximate biomass “reaction”, which specifies biomass composition in terms of precursor metabolites (such as amino acids and nucleotides). On the one hand, biomass composition is often not known exactly and may vary drastically between conditions and strains. On the other hand, the predictions of computational models crucially depend on biomass. Also elementary flux modes (EFMs), which generate the flux cone, depend on the biomass reaction.

To better understand cellular phenotypes across growth conditions, we introduce and analyze new classes of elementary vectors for comprehensive (next-generation) metabolic models, involving explicit synthesis reactions for all macromolecules. Elementary growth modes (EGMs) are given by stoichiometry and generate the growth cone. Unlike EFMs, they are not support-minimal, in general, but cannot be decomposed “without cancellations”. In models with additional (capacity) constraints, elementary growth vectors (EGVs) generate a growth polyhedron and depend also on growth rate. However, EGMs/EGVs do not depend on the biomass composition. In fact, they cover all possible biomass compositions and can be seen as unbiased versions of elementary flux modes/vectors (EFMs/EFVs) used in traditional models.

To relate the new concepts to other branches of theory, we consider autocatalytic sets of reactions. Further, we illustrate our results in a small model of a self-fabricating cell, involving glucose and ammonium uptake, amino acid and lipid synthesis, and the expression of all enzymes and the ribosome itself. In particular, we study the variation of biomass composition as a function of growth rate. In agreement with experimental data, low nitrogen uptake correlates with high carbon (lipid) storage.

**Author summary:** Next-generation, genome-scale metabolic models allow to study the reallocation of cellular resources upon changing environmental conditions, by not only modeling flux distributions, but also expression profiles of the catalyzing proteome. In particular, they do no longer assume a fixed biomass composition. Methods to identify *optimal* solutions in such comprehensive models exist, however, an unbiased understanding of all *feasible* allocations is missing so far. Here we develop new concepts, called elementary growth modes and vectors, that provide a generalized definition of minimal pathways, thereby extending classical elementary flux modes (used in traditional models with a fixed biomass composition). The new concepts provide an understanding of all possible flux distributions *and* of all possible biomass compositions. In other words, elementary growth modes and vectors are the unique functional units in any comprehensive model of cellular growth. As an example, we show that lipid accumulation upon nitrogen starvation is a consequence of resource allocation and does not require active regulation. Our work puts current approaches on a theoretical basis and allows to seamlessly transfer existing workflows (e.g. for the design of cell factories) to next-generation metabolic models.

## 1 Introduction

A major characteristics of life is self-fabrication, involving self-maintenance and self-replication. Cellular self-fabrication requires the acquisition and transformation of nutrients, not only to maintain the cell, but also to replicate, that is, to grow. During one cycle, a cell needs to duplicate all its building blocks and to cover the related costs. In constant environments, this process is balanced and leads to exponential growth. Indeed, exponential growth means that all cellular components are synthesized in proportion to their abundance. Clearly, metabolic activity depends on growth rate since higher growth rates imply higher synthesis rates which in turn require more ribosomes that produce the additional enzymes (and the ribosomes themselves). Thus, growth can be seen as a process that allocates cellular resources, limited by environmental and physico-chemical constraints.

Arguably, some of the most successful approaches to study (microbial) growth processes are rooted in constraint-based modeling [1]. At their heart sits a (genome-scale) metabolic model, which captures all possible (cell-specific) biochemical transformations in an annotated and mathematically structured form. The resulting stoichiometric matrix, coupled with environmental and physico-chemical constraints, is already sufficient to study (fundamental aspects of) growth. Various biased and unbiased methods have been developed to predict (steady-state) metabolic phenotypes [2]. However, first-generation genome-scale metabolic models do not involve the expression of proteins and hence do not account for the individual enzyme costs of metabolic fluxes. These models rather use a fixed biomass composition that represents the average costs of growth. Recent efforts focus on the development of next-generation genome-scale metabolic models that also account for the enzyme demands of individual reactions. As one prominent example, we mention resource balance analysis (RBA) [3]. Although specific approaches differ in their degree of mechanistic detail and mathematical formulation, the central concept in the analysis of next-generation models is the (optimal) allocation of resources. However, we currently lack a theoretical understanding of all feasible allocations in next-generation metabolic models.

In traditional, first-generation models, elementary flux modes and vectors (EFMs [4, 5] and EFVs [6, 7]) allow an unbiased study of metabolic pathways, in particular, an interpretation of any *feasible* flux distribution in terms of unique functional units. On the other hand, flux balance analysis (FBA) enables the computationally efficient identification of *optimal* fluxes [8]. In this respect, RBA can be viewed as a generalization of FBA that also accounts for enzyme costs. It represents the corresponding tool to identify optimal fluxes (and corresponding optimal allocations) in comprehensive, next-generation models. However, the analogues of EFMs and EFVs have not been identified, yet. In [9], “elementary growth states” have been introduced for (semi-)kinetic models of cellular self-fabrication, but for constraint-based models, the corresponding theoretical concepts are still missing.

In this work, we introduce elementary growth modes and vectors (EGMs and EGVs) that are given by stoichiometry and irreversibility (EGMs) as well as by additional constraints and growth rate (EGVs). In fact, we develop a mathematical theory that enables an unbiased characterization of all feasible flux distributions in constraint-based models of cellular growth. In analogy to EFMs and EFVs, EGMs and EGVs can be interpreted as unique functional units of self-fabrication. Even more importantly, they provide an unbiased understanding of all possible biomass compositions. In a small example of a self-fabricating cell, we highlight that the experimentally observed lipid accumulation upon nitrogen starvation is a general feature of a comprehensive, next-generation metabolic model and not the result of active regulation.

Finally, we relate EGMs and EGVs to other branches of theory. Obviously, purely stoichiometric models do not reflect implications from autocatalysis (namely, that all catalysts of active reactions need to be synthesized) and kinetics (namely, that all species involved in active reactions need to be present with nonzero concentrations). We observe that additional (capacity) constraints often ensure that EGVs are autocatalytic.

## 2 Results

In this section, we introduce and analyze *elementary growth modes* and *vectors* for computational models of cellular growth. Further, we relate our approach to the theory of autocatalytic sets. In order to motivate the new concepts and to illustrate our theoretical results, we use a running example (a small model of a self-fabricating cell).

In Section 3 (Methods), we provide the relevant mathematical background (elementary vectors in polyhedral geometry), and in Section 4 (Discussion), we compare our theory with the constraint-based study of traditional models and the analysis of semi-kinetic models [9].

To begin with, we summarize the mathematical notation used throughout this work.

### Notation

We denote the positive real numbers by 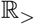 and the nonnegative real numbers by 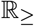. Let *I* be an index set; often *I* = {1, …, *n*}, and 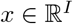 stands for 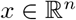. For 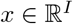, we write *x* > 0 if 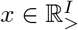, *x* ≥ 0 if 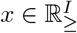, and we denote its *support* by supp(*x*) = {*i* ∈ *I* | *x_i_* ≠ 0}. Recall that a nonzero vector 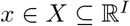 is *support-minimal* (in *X*) if, for all nonzero *x*′ ∈ *X*, supp(*x*′) ⊆ supp(*x*) implies supp(*x*′) = supp(*x*). For 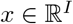, we define its *sign vector* sign(*x*) ∈ {−, 0, +}^*I*^ by applying the sign function component-wise, that is, sign(*x*)_*i*_ = sign(*x_i_*) for *i* ∈ *I*. The relations 0 < – and 0 < + on {−, 0, +} induce a partial order on {−, 0, +}^*I*^: for *X, Y* ∈ {−, 0, +}^*I*^, we write *X* ≤ *Y* if the inequality holds component-wise. For 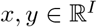, we denote the component-wise product by 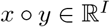, that is, (*x* ∘ *y*)_*i*_ = *x_i_y_i_*. For 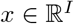 and subindex set 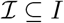, we write 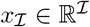 for the corresponding subvector.

### 2.1 Growth model

The dynamic model of cellular growth studied in this work is given by the mass balance equation

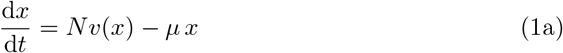

and, additionally, by the (dry) mass constraint

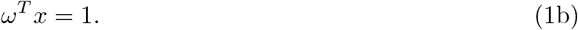

Here, *x* and *ω* are the vectors of concentrations and molar masses (of the molecular species), respectively, *v* is the vector of reaction rates, *N* is the stoichiometric matrix which captures the net effect of the chemical reactions (and other processes), and *μ* is the growth rate. We summarize the fundamental objects and quantities in Table 1 and provide a minimal derivation of the set of Eqs (1) in Supporting Information A. For alternative derivations, see e.g. [10] or [9].

**Table 1.** Fundamental objects and quantities of cellular growth.

| Symbol | Name | Unit |
| --- | --- | --- |
| Mol | set of molecular species | – |
| Rxn | set of chemical reactions | – |
| $N \in \mathbb{R}^{\text{Mol} \times \text{Rxn}}$ | (net) stoichiometric matrix | 1 |
| $\omega \in \mathbb{R}_{>}^{\text{Mol}}$ | molar masses | $\text{g mol}^{-1}$ |
| $x \in \mathbb{R}_{\geq}^{\text{Mol}}$ | concentrations | $\text{mol g}^{-1}$ |
| $v(x) \in \mathbb{R}^{\text{Rxn}}$ | reaction rates | $\text{mol g}^{-1} \text{h}^{-1}$ |
| $\mu \in \mathbb{R}_{\geq}$ | growth rate | $\text{h}^{-1}$ |

In the following, we also consider (dimensionless) mass fractions 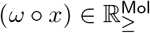. Obviously, 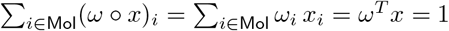.

By Eqs (1), growth rate is given by

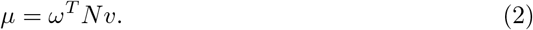

By distinguishing between exchange and internal reactions, Eq (2) can be rewritten as

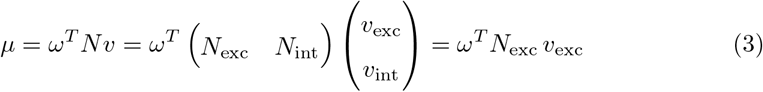

since *ω^T^N*_int_ = 0, by mass conservation. In biological terms, growth rate is determined by the exchange reactions with the environment (the uptake of nutrients and the excretion of waste products).

We highlight two observations, proven in Supporting Information A:

- *Conservation laws*. In a model of cellular growth, there cannot be any conservation laws. In mathematical terms, ker 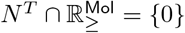.
- *Dependent concentrations*. Still, there can be dependent concentrations. That is, possibly ker *N^T^* ≠ {0}.

In order to motivate the new concepts in this work and to illustrate our results, we will use a running example.

### 2.2 Example

Consider the small model of a self-fabricating cell shown in Fig 1(a). The cell takes up glucose (G) and ammonium (N) via the catalytic reactions

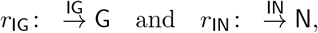

respectively, and forms amino acids (AA) and lipids,

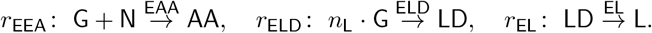

**Fig 1.**
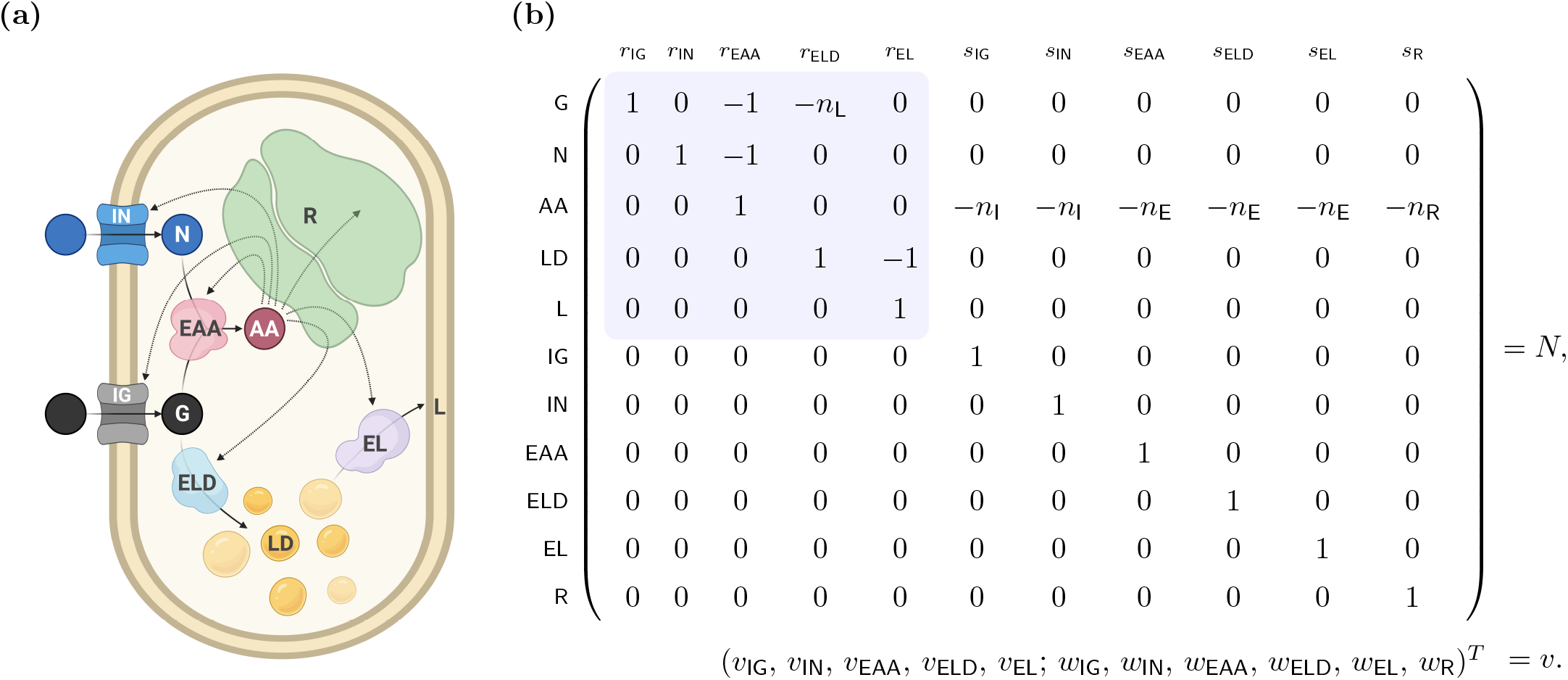
A small model of a self-fabricating cell. **(a)** The cell has two types of importers, taking up glucose (G) and ammonium (N) from the environment, and three types of metabolic enzymes, synthesizing amino acids (AA) from glucose and ammonium, forming lipid droplets (LD) from glucose, and transferring lipids (L) to the membrane. Amino acids are used by the ribosome (R) to synthesize importers (IN, IG), metabolic enzymes (EAA, ELD, EL), and the ribosome itself. **(b)** The resulting stoichiometric matrix and the corresponding flux vector. (Thereby v is used for import and enzymatic reactions and *w* for synthesis reactions.) In traditional stoichiometric models, only the metabolic part of the stochiometric matrix (shaded in blue) is considered (together with a “biomass reaction”), see Fig 8.

In fact, lipids are stored in the form of lipid droplets (LD) and transferred to the membrane (L).

Amino acids are the essential building blocks for the importers (*I* = IG, IN), the enzymes (*E* = EAA, ELD, EL), and the ribosome R,

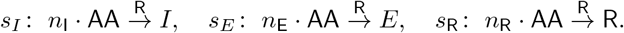

Thereby, stoichiometric coefficients are denoted by *n_i_* with *i* ∈ {L, I, E, R}. The resulting stoichiometric matrix and the corresponding flux vector are displayed in Fig 1(b), and parameter values are given in Table 2.

**Table 2.** Parameter values for the small model of a self-fabricating cell.

| Symbol | Name | Value | Unit | Source* |
| --- | --- | --- | --- | --- |
| $n_L$ | | 7** | 1 | BNID 109367, [21] |
| $n_I$ | | 646 | 1 | MC: CPLX-157 |
| $n_E$ | | 325 | 1 | BNID 108986 |
| $n_R$ | | 22608*** | 1 | BNID 110218 |
| $\omega_G$ | Molar mass glucose | 180 | $\text{g mol}^{-1}$ | |
| $\omega_N$ | Molar mass ammonium | 18 | $\text{g mol}^{-1}$ | |
| $k_{\text{cat}}$ | Turnover number | 79 | $\text{s}^{-1}$ | [22] |
| $k_{\text{el}}$ | Elongation rate | 8 | $\text{AA s}^{-1}$ | BNID 111689 |
| $A_L$ | Lipid area | 0.5 | $\text{nm}^2$ | BNID 106993 |
| $A_I$ | Importer area | 48 | $\text{nm}^2$ | BNID 114685 |
| $r$ | Surface-to-volume ratio | $9.3 - 2.8 \mu\text{h}$ | $\mu\text{m}^{-1}$ | [23] |
| $\rho$ | Cell density | 290 | $\text{g L}^{-1}$ | BNID 104443 |
| $\alpha$ | Minimum lipid fraction | 0.5 | 1 | [21] |

\* For values from BioNumbers [24] and MetaCyc [25] the database IDs are provided (BioNumbers: BNID; MetaCyc: MC).
\*\* Estimated from the number of carbons in a typical *E. coli* lipid.
\*\*\* Number of AA in the ribosome multiplied by three to take into account RNA content.

In the next subsection, we introduce the basic “building blocks” of any possible flux distribution in a (purely stoichiometric) model of cellular growth.

### 2.3 Elementary growth modes

At steady state, Eq (1a) implies

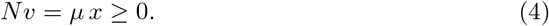

Further, in a given setting, some reactions may have a given direction, as determined by thermodynamics. That is,

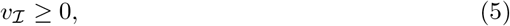

where 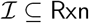 denotes the set of irreversible reactions.

The inequalities *Nv* ≥ 0 and 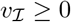 (for the fluxes *v*) specify a polyhedral cone which suggests the following definition.

#### Definition 1.

*Growth modes* (GMs) for the dynamic growth model (1) are elements of the *growth cone*

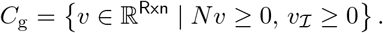

A GM *v* ∈ *C*_g_ has an *associated* growth rate *μ*(*v*) = *ω^T^Nv* ≥ 0 and, if *μ*(*v*) > 0, an *associated* concentration vector 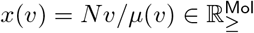.

*Elementary* growth modes (EGMs) are conformally non-decomposable GMs.

At this point, we refer the reader to Subsection 3.1 for an introduction to elementary vectors in polyhedral geometry (in particular, for the definitions of polyhedral cones and conformal non-decomposability). Still, we also try to provide an intuitive understanding of Definition 1:

- GMs are fluxes given by stoichiometry and irreversibility; in particular, they do not depend on concentrations or growth rate. Still, GMs have an associated growth rate and an associated concentration vector, as given by Eqs (2) and (4).
- A growth cone *C*_g_ is a general polyhedral cone. In contrast, a flux cone 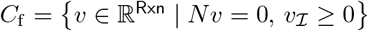, arising also in traditional models of cellular growth, is a special cone lying in a subspace (the nullspace of the stoichiometric matrix). For visualizations of a special and a (more) general polyhedral cone, see Figs 5 and 6 in Subsection 3.1.
- Whereas the elementary vectors of a flux cone are support-minimal, the elementary vectors of a growth cone are conformally non-decomposable. Thereby, a conformally non-decomposable vector cannot be written as a sum of other vectors *without cancellations* [11].
- In applications, the growth cone (as the flux cone) is often pointed (due to irreversibility).

GMs are scalable, but the associated concentrations are scale invariant.

#### Proposition 2.

*For a GM v* ∈ *C*_g_ *with associated concentration x*(*v*) *and* λ > 0, *it holds that x*(λ*v*) = *x*(*v*).

*Proof*.

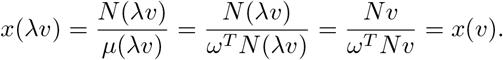

Further, we can characterize the case of zero growth rate.

#### Proposition 3.

*For a GM v* ∈ *C*_g_, *μ*(*v*) =0 *is equivalent to Nv* = 0.

*Proof*. Obviously, *Nv* = 0 implies *μ*(*v*) = *ω^T^Nv* = 0. Conversely, *μ*(*v*) = *ω^T^Nv* = 0 with 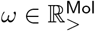 and 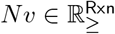 implies *Nv* = 0.

GMs *v* with *μ*(*v*) = 0 are flux modes (FMs), that is, elements of the flux cone

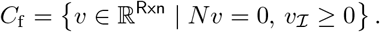

EGMs *v* with *μ*(*v*) = 0 are elementary flux modes (EFMs), that is, support-minimal elements of *C*_f_. They are not support-minimal elements of *C*_g_, in general.

Finally, we apply the general theory of elementary vectors and state the main result of this subsection.

#### Theorem 4.

*Every nonzero GM is a conformal sum of EGMs*.

*Proof*. By Theorem 11 in Subsection 3.1. The growth cone *C*_g_ is a general polyhedral cone, and its elementary vectors are the conformally non-decomposable vectors, that is, the EGMs.

In fact, EGMs with nonzero growth rate can be scaled to have the same growth rate as the given GM.

#### Corollary 5.

*Let v be a nonzero GM with associated growth rate μ*(*v*) ≕ *μ. Then, there exist (possibly empty) finite sets E*_0_ *and E_μ_ of EGMs with associated growth rates* 0 *and μ* > 0, *respectively, such that*

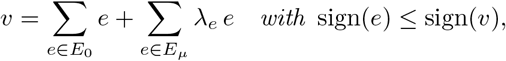

λ*_e_* ≥ 0, *and* 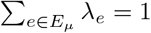. *Moreover, if μ* > 0, *then* 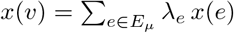.

*Proof*. By Theorem 4, there exist (possibly empty) finite sets *E*_0_ and *E*_>_ of EGMs (with associated growth rates 0 and *>* 0, respectively) such that

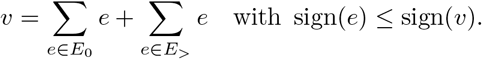

In particular, 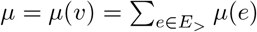. If *μ* > 0, that is, 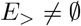, then

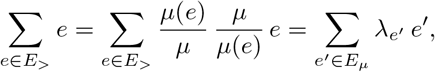

where 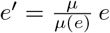 with *μ*(*e*′) = *μ* and 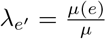 with 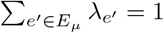.

Finally, if *μ* > 0, then 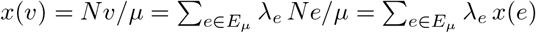

In Corollary 5, we actually fix growth rate which turns the growth cone into a polyhedron. Hence, the result is also an instance of Theorem 12 in Subsection 3.1.

#### Example (EGMs)

Recall the small model of a self-fabricating cell given in Fig 1 and note that all reactions are assumed to be irreversible, 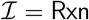. As it turns out, there are 11 EGMs (up to scaling), corresponding to the 11 molecular species, that is, 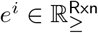 with *i* ∈ Mol = {G, N, AA, LD, L, IG, IN, EAA, ELD, EL, R}. Explicitly,

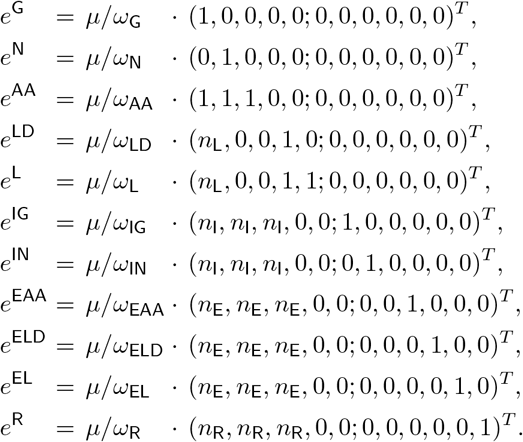

Thereby, we introduced the factors *μ/ω_i_* to obtain correct units (molg^−1^ h^−1^) and equal growth rate *μ* for all EGMs. We note the following points:

- Every EGM “produces” exactly one molecular species, as indicated by its name. (For every EGM, there is exactly one molecular species with nonzero associated concentration.) Formally, *Ne^i^* = (*μ*/*ω_i_*)*u^i^*, where *u^i^* is the *i*th unit vector in 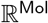. That is, *e*^G^ produces G (glucose), *e*^N^ produces N (ammonium), …, and *e*^R^ produces R (ribosome). This special situation arises from the fact that the stoichiometric matrix *N* is square. In general, there may be more than one EGM for the exclusive production of some molecular species or, conversely, no EGM for exclusive production (just joint production with another species).
- EGMs *e*^G^ and *e*^N^ are support-minimal, but all other EGMs are not.
- For the associated growth rates, we obtain 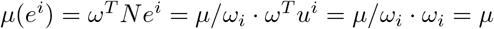. (This is just the consequence of introducing the factors above.) For the associated concentrations, we obtain 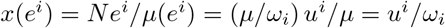.
- There are no EGMs with zero growth rate, that is, there are no EFMs.

To give an example of a conformal sum, we consider the GM

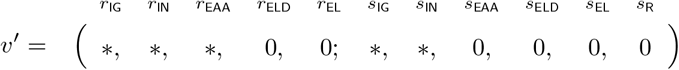

with support in the import/enzymatic reactions *r*_IG_, *r*_IN_, *r*_EAA_ and in the synthesis reactions *s*_IG_, *s*_IN_. In particular, *v*′ is a GM without lipid synthesis. Now, let *μ*(*v*′) = *μ*. By Theorem 4 (and Corollary 5), *v*′ is a conformal (and convex) sum of EGVs,

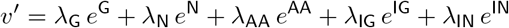

with λ_G_, λ_N_, λ_AA_ ≥ 0, λ_IG_, λ_IN_ > 0 and λ_G_ + λ_N_ + λ_AA_ + λ_IG_ + λ_IN_ = 1. Note that *v*′ produces IG, IN (since λ_IG_, λ_IN_ > 0), however, it produces G, N, AA if and only if also λ_G_, λ_N_, λ_AA_ > 0.

Since GMs are given by stoichiometry (and irreversibility), they do not reflect constraints implied by autocatalysis and kinetics.

#### Example (autocatalysis and kinetics)

Obviously, the GM *v*′ above involves catalytic reactions. In particular, it involves *r*_EAA_ (amino acid synthesis) as well as *s*_IG_, *s*_IN_ (the expression of the importers). However, it is not catalytically closed in the sense that reactions *r*_EAA_ and *s*_IG_, *s*_IN_ carry fluxes, but the corresponding catalysts, the enzyme EAA and the ribosome R, are not expressed. Now, EGV *e*^EAA^ involves *s*_EAA_ (the expression of EAA), and analogously *e*^R^ involves *s*_R_. Hence, we form a convex sum

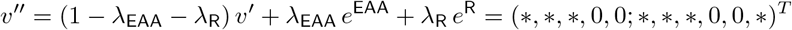

with λ_EAA_, λ_R_ > 0, to obtain a GM that is catalytically closed.

Further, the EGM *e*^G^ is kinetically consistent in the sense that it involves the reaction *r*_IG_ and hence the species G *and* it has a corresponding nonzero associated concentration *x*_G_(*e*^G^) > 0, as implied by kinetics. By the discussion above, the GM *v*′ is kinetically consistent if and only if λ_G_, λ_N_, λ_AA_ > 0. In this case, all species involved in active reactions have nonzero associated concentrations.

In the next subsection, we elaborate on the concepts of catalytic closure and kinetic consistency.

### 2.4 Implications from autocatalysis and kinetics

Cellular growth is autocatalytic in the sense that the cell fabricates itself (thereby exchanging substrates/products with the environment). One needs to distinguish this notion of “network autocatalysis” from “autocatalytic subnetworks” [12, 13].

Consider the overall reaction corresponding to (the flux through) a subnetwork. If a molecular species appears on both the educt and product sides, in particular, with a larger stoichiometric coefficient on the product side than on the educt side, then it is *formally autocatalytic* (cf. [14]). In fact, there are several competing notions of autocatalytic species and subnetworks (cf. [12–14]).

In this work, we consider network autocatalysis. Before we state possible definitions, we distinguish two modeling approaches.

- **Detailed models** (without individual catalytic reactions): In this approach, catalysis occurs on the level of (small) subnetworks. In particular, individual reactions are not catalytic. For example, a simple catalytic mechanism (involving enzyme E, substrate S, and product P) is given by E + S ↔ ES ↔ EP ↔ E + P.
- **Coarse-grained models** (with individual catalytic reactions): In this approach, catalysis occurs on the level of individual reactions. For example, the catalytic mechanism above is written as E + S ↔ E + P or 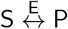. Due to coarse-graining, catalysis cannot be identified from the stoichiometric matrix. Hence, for every catalytic reaction, the corresponding catalyst is specified explicitly.

For **detailed models**, one may call a growth mode autocatalytic if it contains an autocatalytic species or subnetwork and it is catalytically closed. Formal definitions and their comparison are beyond the scope of this work. For **coarse-grained models** (like the small model of a self-fabricating given in Fig 1), we give a formal definition of network autocatalysis.

#### Definition 6.

For a coarse-grained model, let Cat ⊆ Rxn be the set of catalytic reactions. A GM *v* ∈ *C*_g_ is *basically catalytic* (BC) if there is a catalytic reaction *r* ∈ supp(*v*) ⋂ Cat. Further, a GM *v* ∈ *C*_g_ is *catalytically closed* (CC) if, for every catalytic reaction *r* ∈ supp(*v*) ⋂ Cat, it holds that (*Nv*)_*s*_ > 0 for the corresponding catalyst *s* ∈ Mol. Finally, a GM *v* ∈ *C*_g_ is *autocatalytic* (AC) if it is BC and CC.

A subset of reactions *S* ⊆ Rxn is *autocatalytic* (AC) if there exists an autocatalytic GM *v* ∈ *C*_g_ with *S* = supp(*v*). A nonempty subset of reactions is *minimally* autocatalytic (MAC) if it is AC and inclusion-minimal.

In the literature, a closure condition is also crucial in the definitions of “reflexive autocatalysis” [15–17] and “chemical organizations” [18–20].

We note the following points:

- AC is implied by two conditions: BC guarantees that there is at least one active catalytic reaction, and CC ensures that all active catalysts are produced. For an illustration, recall the motivating paragraph “Example (autocatalysis and kinetics)” just before this subsection. In the running example, all reactions are catalytic, and hence all GMs are BC. Whereas the GM *v*′ is not CC (the active enzyme EAA and the ribosome R are not produced), the GM *v*″ is CC and hence AC. Its support supp(*v*″) = {*r*_IG_, *r*_IN_, *r*_EAA_; *s*_IG_, *s*_IN_, *s*_EAA_, *s*_R_} is an AC subset of reactions; in fact, it is the only MAC subset of reactions.
- Catalytic closure is defined for fluxes, but it refers to concentrations via *x*(*v*)_*s*_ = (*Nv*)_*s*_/*μ*(*v*). In particular, (*Nv*)_*s*_ > 0 implies *x*(*v*)_*s*_ > 0.

In addition to autocatalysis, also kinetics implies constraints on growth modes. In particular, a growth mode is kinetically consistent if all species involved in active reactions (not necessarily all species in the model) have nonzero associated concentrations.

#### Definition 7.

A GM *v* ∈ *C*_g_ is *kinetically consistent* if, for every *r* ∈ supp(*v*) ⊆ Rxn and *s* ∈ Mol, *N_sr_* ≠ 0 implies (*Nv*)_*s*_ > 0.

We note the following points:

- Like catalytic closure, kinetic consistency is given by stoichiometry.
- Interestingly, kinetic consistency implies formal autocatalysis for all (not necessarily catalytic) species involved.

In the next subsection, we consider additional (capacity) constraints which often ensure catalytic closure.

### 2.5 Constraint-based models

For many systems, kinetic models are not yet available, and constraint-based models are used. Steady-state reaction rates (fluxes) *v* are considered as independent variables, that is, the non-linear dependence of the kinetics on the concentrations *x* is neglected.

Most importantly, catalytic processes imply linear capacity constraints for *x* and *v*. Additional constraints can be formulated for processes that are not catalytic (in the given model), e.g. lower bounds for concentrations or fluxes. In compact form, linear constraints can be can be written as

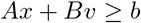

with 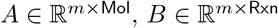, and 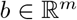.

Altogether, *constraint-based* growth models involve steady-state, irreversibility, (dry) mass, and additional linear constraints,

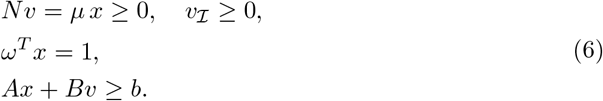

#### Example (additional constraints)

Recall the small model of a self-fabricating cell given in Fig 1. In addition to steady-state, irreversibility, and (dry) mass constraints, we consider capacity constraints for all catalysts (importers, enzymes, and the ribosome) and membrane constraints.

Let *k*_cat_ be the turnover number of the importers IG, IN and enzymes EAA, ELD, EL and *k*_el_ be the elongation rate of the ribosome R. The resulting capacity constraints are given by

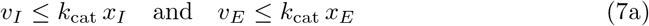

for *I* = IG, IN and *E* = EAA, ELD, EL and

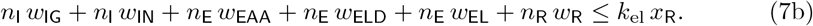

The cell membrane area is formed by lipids L and importers IG and IN, leading to the (equality) constraint

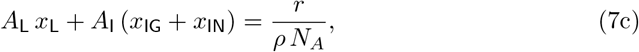

where *A*_L_ and *A*_I_ denote the areas of lipids and importers, respectively, *r* denotes the surface-to-volume ratio, *ρ* denotes cell density, and *N_A_* is Avogadro’s number.

Additionally, we require that a minimum fraction *α* of the surface area is formed by lipids, leading to the (inequality) constraint

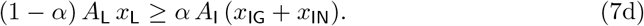

For the derivation of the membrane constraints, see Supporting Information B. As stated above, the additional constraints (7) can be summarized as *Ax* + *Bv* ≥ *b*.

All constraints are based on realistic data. The parameter values for the small model of a self-fabricating cell are given in Table 2.

In the final subsection, we introduce the basic “building blocks” of any possible flux distribution in a constraint-based model of cellular growth.

### 2.6 Elementary growth vectors

For given *μ* > 0 and flux vector *v*, the concentration vector *x* = *Nv/μ* is not an independent variable. In particular, we have *Nv* = *μx* ≥ 0 and *ω^T^Nv* = *ω^T^*(*μx*) = *μ*. Hence, the constraint-based growth model (6) is equivalent to constraints just in terms of *v*,

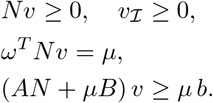

The inequalities above (for the fluxes *v*) specify a polyhedron which suggests the following definition.

#### Definition 8.

Let *μ* > 0. *Growth vectors* (GVs) for the constraint-based growth model (6) are elements of the *growth polyhedron*

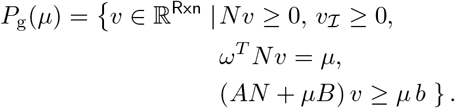

A GV *v* ∈ *P*_g_(*μ*) has an *associated* concentration vector 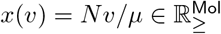.

*Elementary* growth vectors (EGVs) are convex-conformally non-decomposable GVs and conformally non-decomposable elements of the recession cone

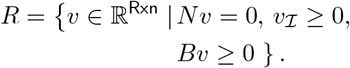

For the definitions of polyhedra, convex-conformal non-decomposability, and the recession cone, we refer the reader to Subsection 3.1. Still, we also try to provide an intuitive understanding of Definition 8:

- GVs are fluxes given by stoichiometry, irreversibility, additional constraints, *and* growth rate. (If growth rate was not fixed, then the problem would become nonlinear, and the resulting set would not be a polyhedron.) In particular, GVs do not depend on concentrations. Still, GVs have an associated concentration vector.
- Every polyhedron can be written as the sum of a polytope and the recession cone. Moreover, every polyhedron has a unique set of conformal generators, and these are the convex-conformally non-decomposable vectors (generating a polytope) and the conformally non-decomposable elements of the recession cone. Thereby, a convex-conformally non-decomposable vector cannot be written as a convex sum of other vectors without cancellations [11].
- The recession cone of the growth polyhedron is a subset of the flux cone. In applications, the recession cone (as a subset of the flux cone) is often pointed (due to irreversibility).

EGMs and EGVs can be computed directly, by algorithms for vertex enumeration (such as `lrs` [26]), or indirectly (after transforming a system of inequalities to a system of equalities), by algorithms for EFM enumeration (such as `efmtool` [27]).

Again, we apply the general theory of elementary vectors and state the main result of this subsection.

#### Theorem 9.

*Let v be a GV for growth rate μ* > 0. *Then, there exist finite sets E*_0_ ⊆ *R and E_μ_* ⊆ *P*_g_(*μ*) *of EGVs such that*

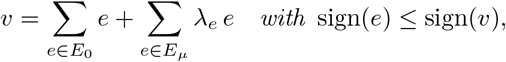

λ_*e*_ ≥ 0, and 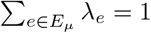. *Moreover*, 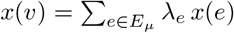.

*Proof*. By Theorem 12 in Subsection 3.1. The growth polyhedron *P*_g_(*μ*) is a general polyhedron, and its elementary vectors are the convex-conformally non-decomposable vectors and the conformally non-decomposable vectors of its recession cone.

Note that every GV is also a GM, and hence every nonzero GV is also a conformal sum of EGMs.

#### Example (EGVs)

Recall the small model of a self-fabricating cell given in Fig 1 and consider the additional constraints (7). For convenience, the essential constraints are summarized in Table S1 in Supporting Information C. We compute EGVs as functions of growth rate, and we observe two distinct regimes. From zero to a critical growth rate *μ*_crit_ = 1.21 h^−1^, we find 24 EGVs. From *μ*_crit_ to the maximum growth rate *μ*_max_ = 1.27 h^−1^, we find 10 EGMs. Eight of these EGVs exist for all growth rates, 16 only in regime L (0 < *μ* < *μ*_crit_) and two only in regime H (*μ*_crit_ < *μ* < *μ*_max_). At *μ*_crit_, two sets of eight EGVs (existing in regime L) merge into two EGVs (in regime H). At *μ*_max_, all EGVs (existing in regime H) merge. EGVs and their characteristic properties are summarized in Table 3.

**Table 3.**
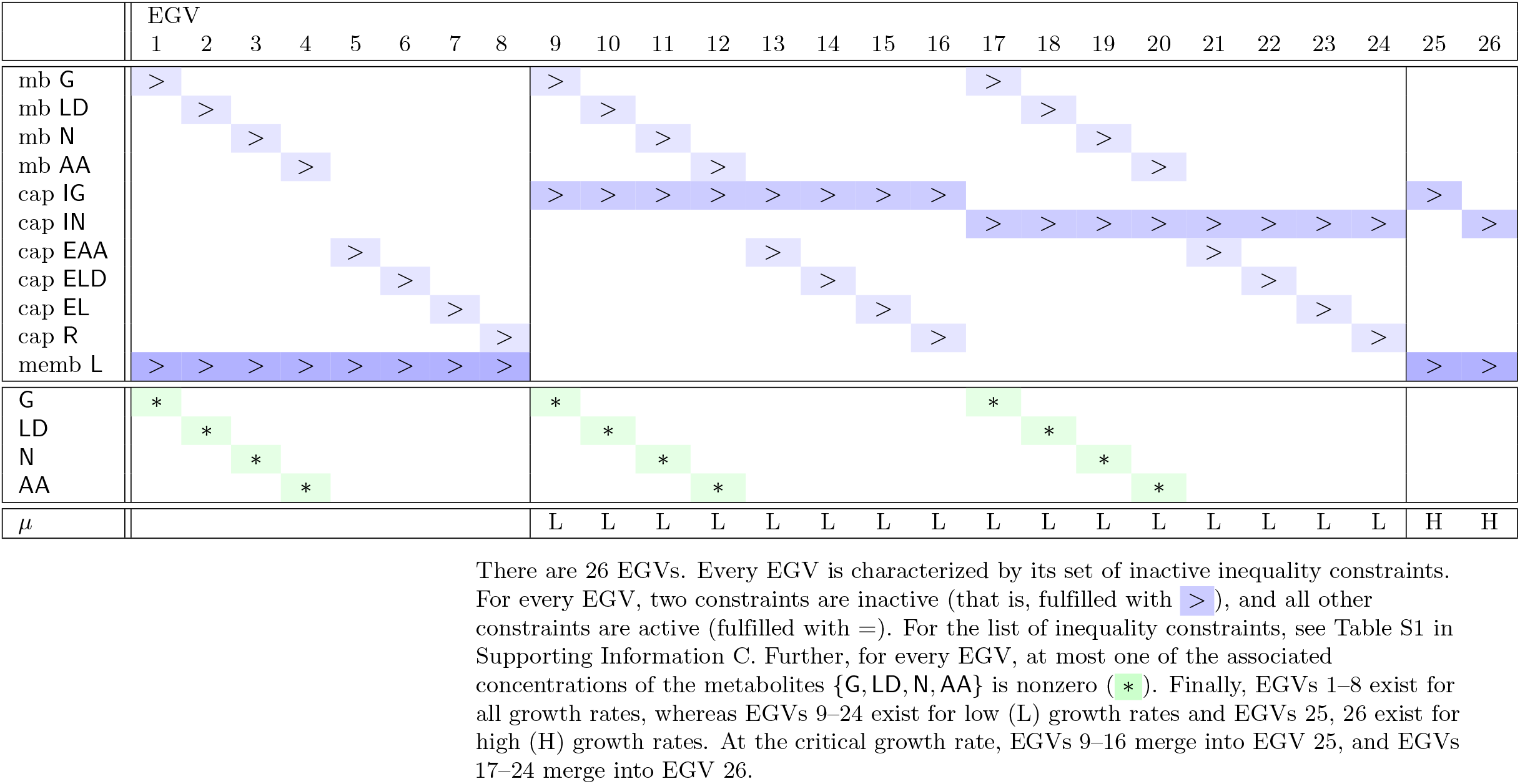
EGVs for the small model of a self-fabricating cell.

Totally, there are 26 EGVs, and every EGV has full support. This is a consequence of the additional (capacity and membrane) constraints which ensure that all reactions are active. As a consequence, every EGV is autocatalytic, and the set of all reactions is the only autocatalytic set of reactions.

In contrast to EFMs, EGVs are not determined by their supports, but they are *characterized* by their sets of *inactive* inequality constraints. For every EGV, two constraints are inactive, and all other constraints are active. Further, for every EGV, the associated concentrations of lipids L and macromolecules {IG, IN, EAA, ELD, EL, R} are nonzero. However, at most one of the associated concentrations of the metabolites {G, LD, N, AA} is nonzero. That is, no EGV is kinetically consistent.

In the example, the cell takes up only glucose and ammonium (and does not excrete any species). Hence, Eq (3) takes the form *μ* = *ω*_G_ *v*_IG_ + *ω*_N_ *v*_IN_, and we introduce the relative contributions of the import fluxes to growth rate,

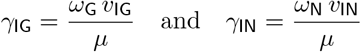

with *γ*_IG_ + *γ*_IN_ = 1. The relative contribution of ammonium uptake to growth as a function of growth rate is visualized in Fig 2.

**Fig 2.**
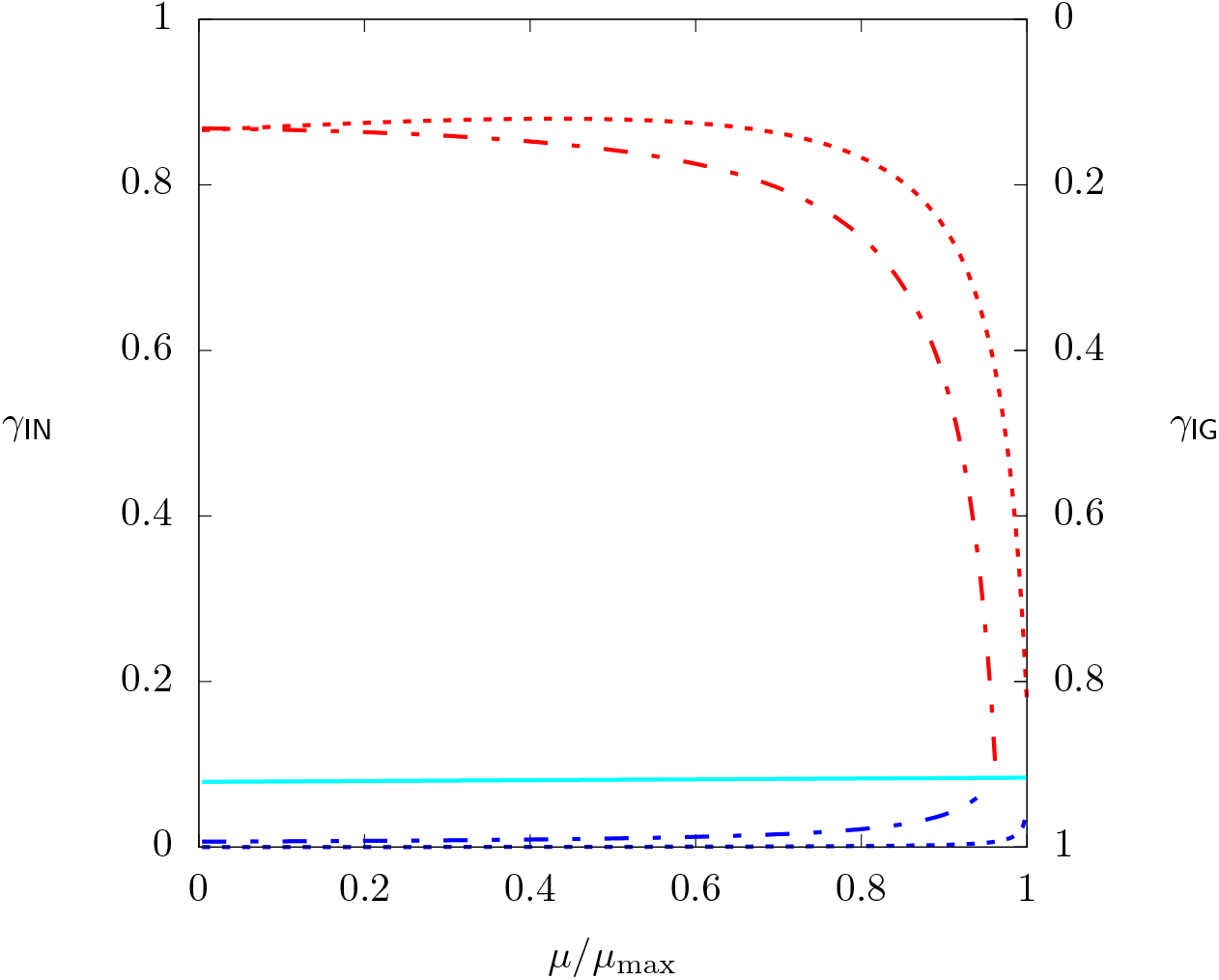
Relative contribution of ammonium uptake to growth, 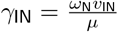, as a function of growth rate for all EGVs (for the small model of a self-fabricating cell). Note that *γ*_IG_ = 1 – *γ*_IN_. Colors correspond to classes of EGVs described in the main text.

As it turns out, EGVs can be grouped into six classes regarding their uptake/storage behavior, namely into classes

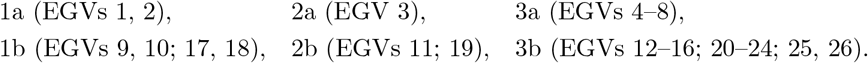

Thereby, EGVs in classes 1a, 2a, and 3a exist for all growth rates, whereas EGVs in classes 1b, 2b, and 3b exist either in regime L or H.

Fig 2 highlights the uptake behaviors of the classes of EGVs. In classes 3a and 3b (turquoise), we observe an (almost) constant ammonium uptake of *γ*_IN_ = 0.091 (independently of growth rate). This corresponds to equal uptake rates of glucose and ammonium. In classes 1a (blue, dotted) and 1b (blue, dashed), ammonium uptake rate is smaller than glucose uptake rate, whereas in classes 2a (red, dotted) and 2b (red, dashed), it is larger. However, all EGVs approach balanced uptake for high growth rates. Classes 1b and 2b (dashed, existing in regime L) reach balanced uptake at *μ*_crit_, whereas classes 1a and 2a (dotted, existing for all growth rates) reach it at *μ*_max_.

For classes with unbalanced uptake behavior, there is an asymmetry between ammonium and glucose uptake at *μ* → 0. For smaller ammonium uptake, *γ*_IN_ → 0, whereas for larger ammonium uptake *γ*_IN_ → 0.87 (instead of *γ*_IN_ → 1). This asymmetry is related to the membrane equality constraint (7c).

Next, we analyze the correlation between ammonium uptake *γ*_IN_ and the mass fraction *s*_C_ = *ω*_G_ *x*_G_ + *ω*_LD_ *x*_LD_ + *ω*_L_ *x*_L_, corresponding to carbon storage in glucose and, most importantly, in lipids. As expected, when ammonium is limiting, then glucose is accumulated or rerouted towards lipid synthesis (instead of amino acid synthesis). Ammonium uptake *γ*_IN_ and carbon storage *s*_C_ as a function of growth rate are visualized in Fig 3.

**Fig 3.**
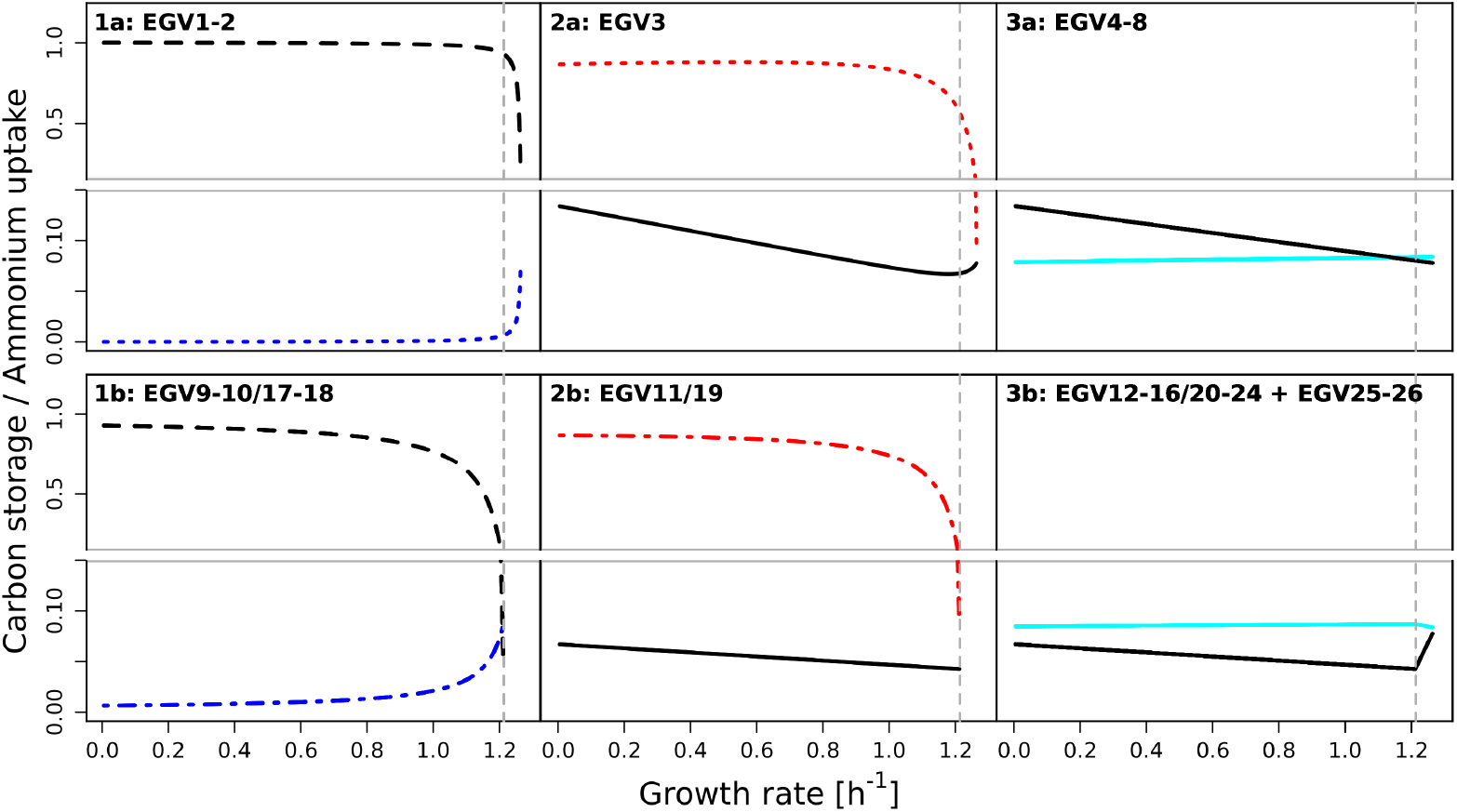
Ammonium uptake 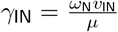 and carbon storage *s*_C_ = *ω*_G_*x*_G_ + *ω*_LD_*x*_LD_ + *ω*_L_*x*_L_ as a function of growth rate for all EGVs (for the small model of a self-fabricating cell). Panels correspond to classes of EGVs described in the main text. Ammonium uptake in blue/red/turquoise (see also Fig 2). Carbon storage in black (dashed and solid: *x*_L_ > 0; dashed: *x*_G_ = 0, *x*_LD_ > 0 or *x*_G_ > 0, *x*_LD_ = 0; solid: *x*_G_ = *x*_LD_ = 0.)

In particular, Fig 3 highlights the uptake/storage behaviors of the six classes of EGVs defined above. The left panels correspond to EGVs in classes 1a and 1b. For low growth rate, they display low ammonium uptake and high carbon storage. The middle panels correspond to classes 2a and 2b, where EGVs display high ammonium uptake and low carbon storage. In fact, ammonium is accumulated. Finally, the right panels correspond to classes 3a and 3b. EGVs display balanced uptake behavior and low carbon storage.

In Supporting Information C, we provide details about the individual EGVs. For each of the 26 EGVs, we display the fluxes (of the 11 reactions) and the associated mass fractions (of the 11 species), as a function of growth rate. See Figs S1 and S2.

Conversely, for each of the 11 species, we show their mass fractions in the 26 EGVs, and analogously, for each of the 11 reactions, we show their fluxes in the 26 EGVs. See Fig 4 in the main text and Fig S3 in Supporting Information C.

**Fig 4.**
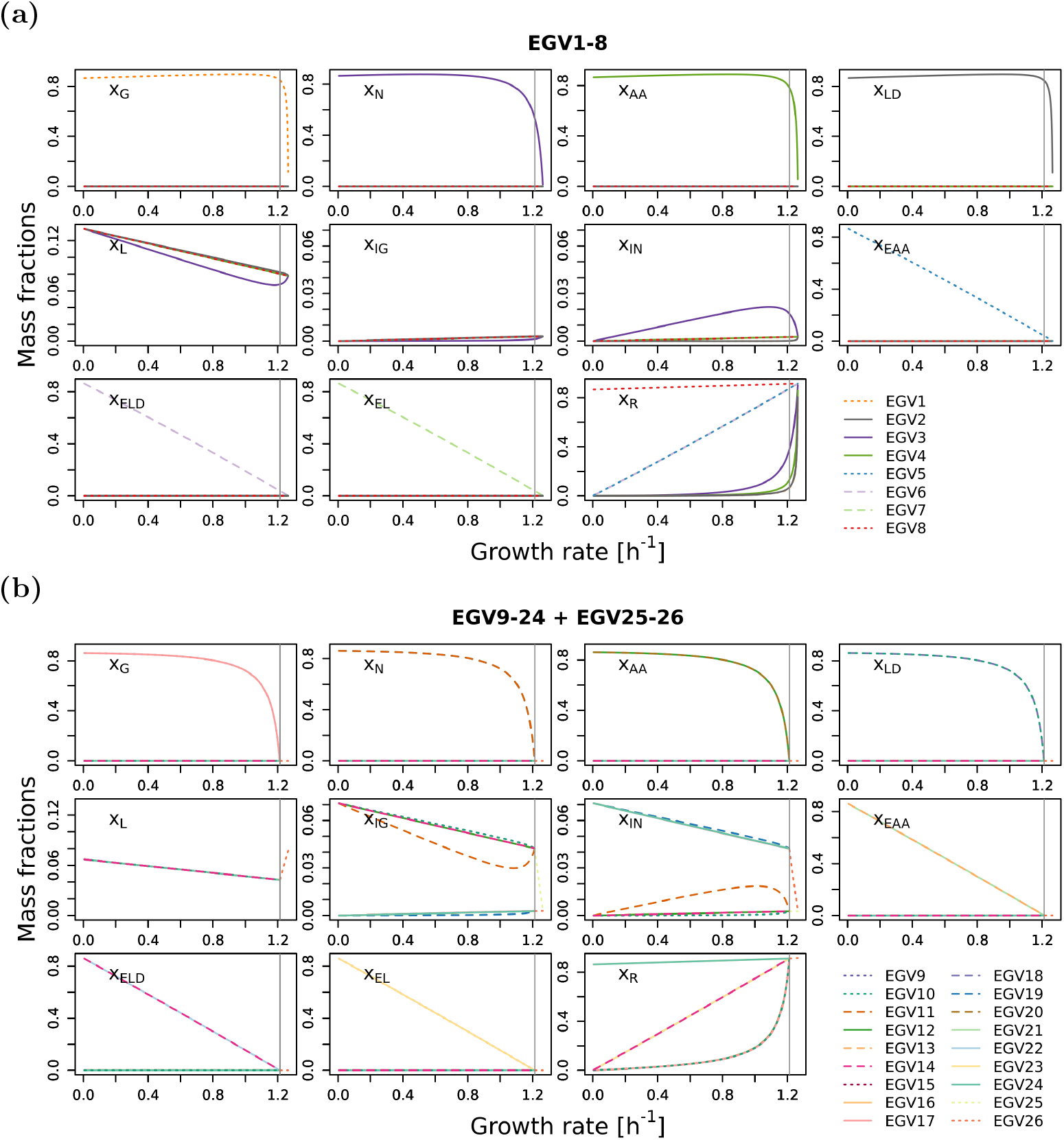
Mass fractions for (a) the 8 EGVs that exist for all growth rates and (b) the 16 EGVs that exist in regime L plus the 2 EGVs that exist in regime H.

### 2.7 Biological interpretation

We used our theoretical concepts to study the storage behavior in the small model of a self-fabricating cell given in Fig 1. In particular, we identified 26 EGVs in three classes regarding their storage behaviors (cf. Fig 3):

1. carbon accumulates in the form of glucose or lipids;
2. nitrogen accumulates in the form of ammonium;
3. ammonium and glucose are stored together in the form of proteins.

We observed that carbon accumulation (class 1) goes hand in hand with low ammonium uptake. In fact, a common strategy to increase cellular carbon content in biotechnological applications is to limit the source of nitrogen [28–36]. Restricting nitrogen limits the synthesis of amino acids and proteins, so the remaining carbon source can be rerouted towards lipid and carbohydrate synthesis, which does not require nitrogen. Remarkably, this effect is a feature of a constraint-based metabolic model and does not require active regulation. Experimentally, this was also observed for an oleaginous fungus, where nitrogen limitation leads to triglyceride accumulation without an up-regulation of the triglyceride synthesis genes, which suggests that it results from a reallocation of resources [37]. Experimentally, nitrogen limitation sometimes leads to the accumulation of fat [28–36] or starch with [32, 34] or without [38] a simultaneous lipid accumulation. All of these possibilities are covered by EGVs and agree with our observations. Of course, which of these behaviors is realized is the result of cellular regulation and hence outside the scope of our analysis.

Further, nitrogen accumulation (class 2) takes place when ammonium uptake exceeds glucose import. In fact, most of the ammonium accumulates since protein synthesis is limited by insufficient glucose import. Still, carbohydrate concentrations remain at normal levels. Such phenotypes have been observed in species such as diatoms, foraminifers, and fungi [39].

The last behavior (class 3) occurs when ammonium and glucose uptake are balanced. The individual EGVs correspond to situations where a certain protein’s abundance is larger than required by the flux level of the catalyzed reaction [40, 41]. This mirrors the fact that enzyme expression is not necessarily indicative of flux levels.

At high growth rate, storage becomes exceedingly expensive for a cell since most resources are required to support growth [40]. Consistently, all EGVs show decreasing storage with increasing growth rate. At maximum growth rate, all EGVs (that exist for high growth rate) merge. Uptake is balanced, and there is no storage (except lipids in the membrane).

Alternatively, the 26 EGVs can be grouped into three classes regarding their ribosome content, namely into classes

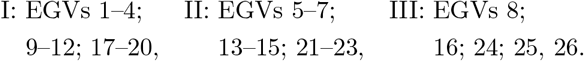

Fig 5 highlights the ribosome mass fraction *ω*_R_ *x*_R_ (as a function of growth rate) of the three classes.

**Fig 5.**
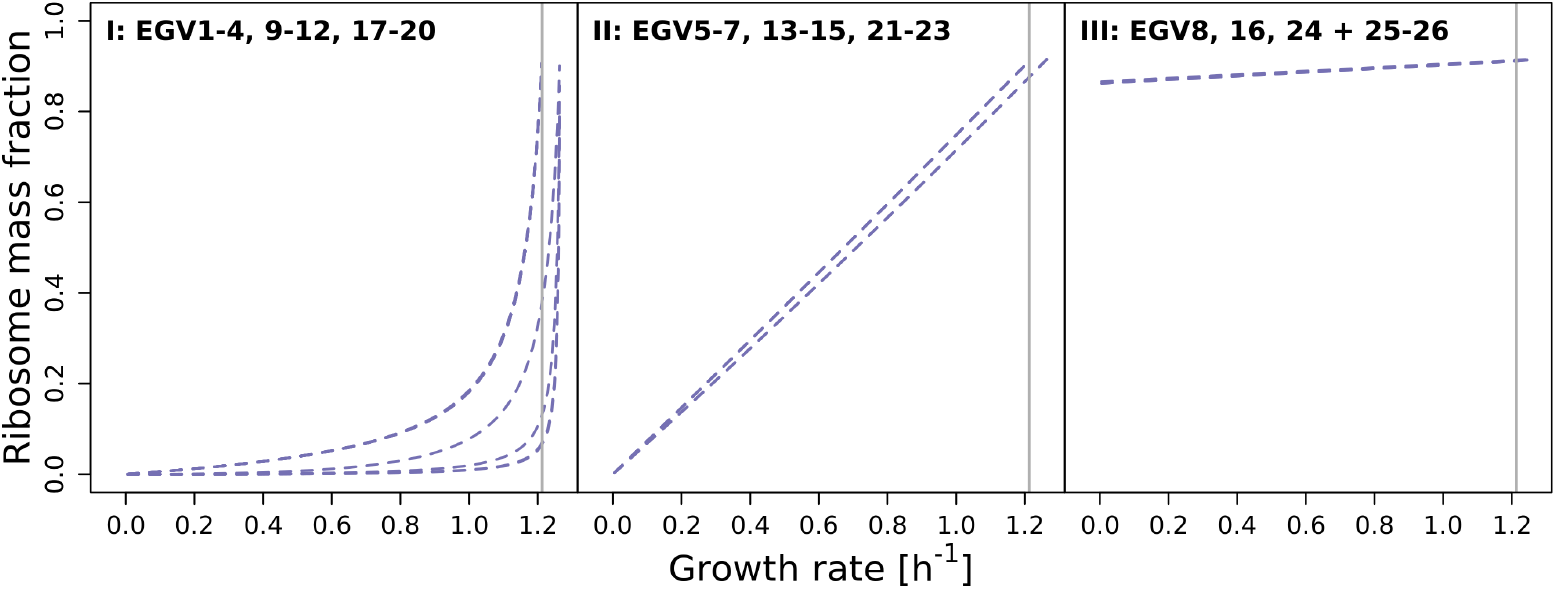
Ribosome mass fraction as a function of growth rate for all EGVs (for the small model of a self-fabricating cell). Panels correspond to classes of EGVs described in the main text.

In class II, the ribosome content is linear as a function of growth rate. This behavior has been observed experimentally (however, with an offset at zero growth rate) [42] and explained theoretically (thereby assuming growth rate-independent “housekeeping” proteins) [43].

In class III, we observe an (almost) constant (and high) ribosome mass fraction of *ω*_R_ *x*_R_ = 0.9 (independently of growth rate). EGVs in this class may describe exactly those ribosomes that are required for the production of growth rate-independent “housekeeping” proteins (and explain the offset at zero growth rate).

In class I, the ribosome content is non-linear at high growth rates. EGVs in this class have nonzero associated metabolite concentrations and may explain a potential non-linear behavior.

Altogether, the experimentally observed ribosome mass fraction [42] can be understood as a convex, conformal sum of EGVs from the three classes.

## 3 Methods

### 3.1 Elementary vectors in polyhedral geometry

For the objects of polyhedral geometry (subspaces, cones, polyhedra), there is no *unique* minimal set of generators, in general. However, *elementary vectors* (EVs) form unique sets of *conformal* generators [11, Section 3.4]. For linear subspaces and s-cones (arising from linear subspaces and nonnegativity constraints), elementary vectors are the support-minimal (SM) vectors; for general polyhedral cones, they are the conformally non-decomposable (cND) vectors; and for general polyhedra, they are the convex-conformally non-decomposable (ccND) vectors plus the cND vectors of the recession cone [11].

Below, we summarize basic definitions and results for s-cones, general polyhedral cones, and polyhedra.

#### S-cones

Given a linear subspace 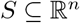 and a subindex set 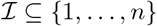, an *s-cone* (special cone, subspace cone) is given by 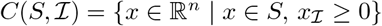. Note that a linear subspace is an s-cone, 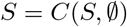.

A vector 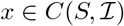 is *elementary* if it is SM. (For linear subspaces, the definition of elementary vectors (EVs) as SM vectors was given in [44].)

The following result is fundamental. See [11, Theorem 3] based on [44, Theorem 1].

##### Theorem 10.

*Let* 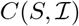 *be an s-cone. Every nonzero vector* 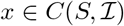 *is a conformal sum of EVs. That is, there exists a finite set E of EVs such that*

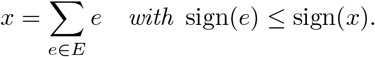

*The set E can be chosen such that* |*E*| ≤ *dim*(*S*) *and* |*E*| ≤ |supp(*x*)|.

##### Example

Consider a minimal metabolic network, involving one internal molecular species *X*, where substrate is taken up, and two products are excreted, that is, (*S*) → *X*, *X* → (*P*_1_), *X* → (*P*_2_), leading to the stoichiometric matrix

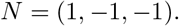

The flux cone 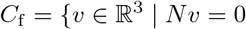 and *v* ≥ 0} is an s-cone, 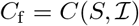 with *S* = ker *N* and 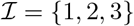 (since all reactions are assumed to be irreversible). The cone is generated by two (support-minimal) EVs, *e*^2^ = (1, 1, 0)^*T*^ and *e*^3^ = (1, 0, 1)^*T*^. The cone is indicated in Fig 6.

**Fig 6.**
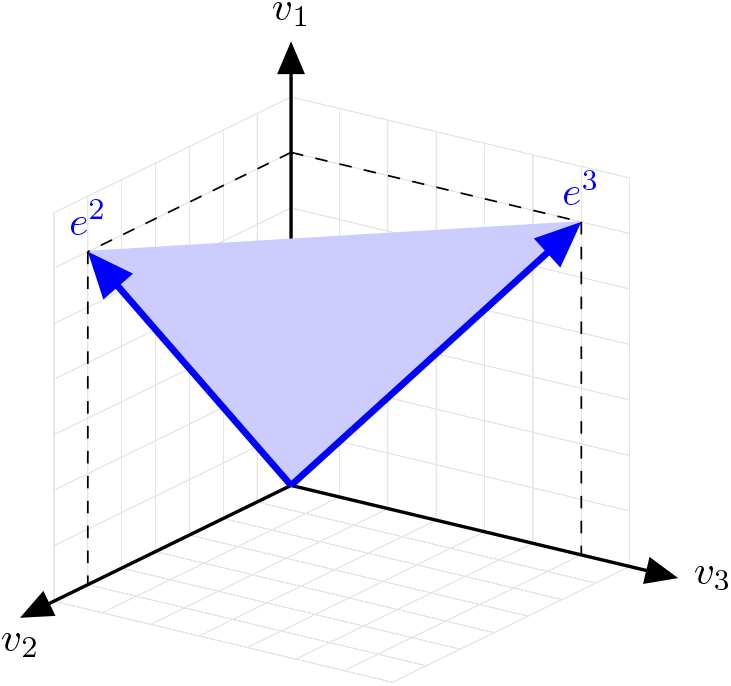
An s-cone.

#### General polyhedral cones

Let *C* be a polyhedral cone, that is, 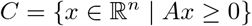 for some 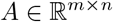. A nonzero vector *x* ∈ *C* is *conformally non-decomposable* (cND) if, for all nonzero *x*^1^, *x*^2^ ∈ *C* with sign(*x*^1^), sign(*x*^2^) ≤ sign(*x*), the decomposition *x* = *x*^1^ + *x*^2^ implies *x*^1^ = λ*x*^2^ with λ > 0. A vector *x* ∈ *C* is *elementary* if it is cND.

By defining EVs as cND vectors (instead of SM vectors), Theorem 10 can be extended to general polyhedral cones. See [11, Theorem 8].

##### Theorem 11.

*Let* 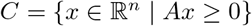 *be a polyhedral cone. Every nonzero vector x* ∈ *C is a conformal sum of EVs. That is, there exists a finite set E of EVs such that*

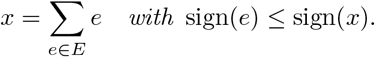

*The set E can be chosen such that* |*E*| ≤ dim(*C*) *and* |*E*| ≤ | supp(*x*)| + | supp(*Ax*)|.

##### Example

Consider a minimal growth model, involving three internal molecular species, a precursor *P*, a metabolic “enzyme” *E*, and the ribosome *R*, where the syntheses of *P*, *E*, and *R* are catalyzed, that is, 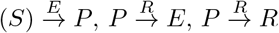, leading to the stoichiometric matrix

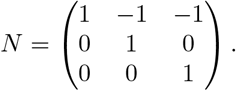

The growth cone 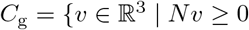 and *v* ≥ 0} is a general polyhedral cone, that is, 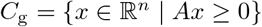 with 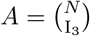 where I_3_ is the identity matrix. The cone is generated by three (conformally non-decomposable) EVs, *e*^1^ = (1, 0, 0)^*T*^, *e*^2^ = (1, 1, 0)^*T*^, and *e*^3^ = (1, 0, 1)^*T*^. (Note that only *e*^1^ is SM.) The cone is indicated in Figure 7.

**Fig 7.**
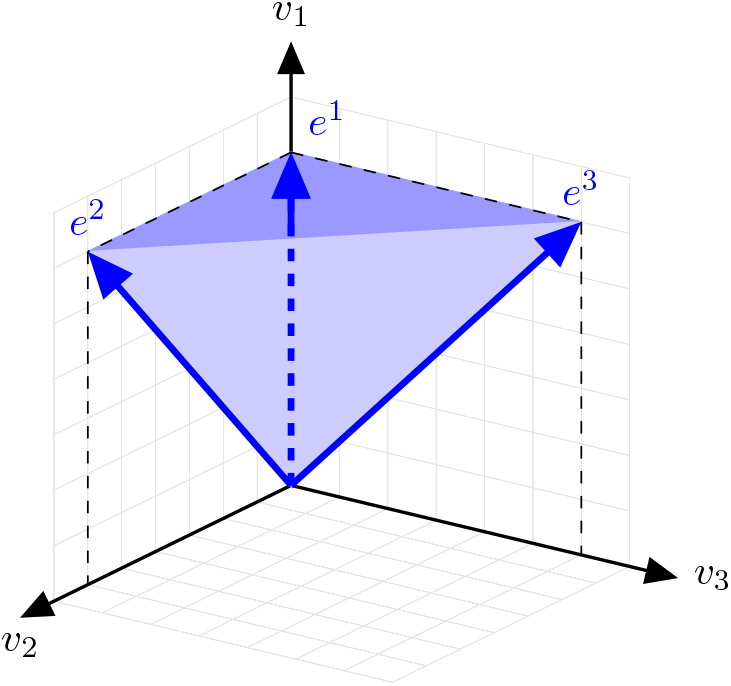
A polyhedral cone (that is not an s-cone).

#### Polyhedra

Let *P* be a polyhedron, that is, 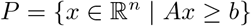 for some 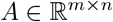 and 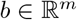. A vector *x* ∈ *P* is *convex-conformally non-decomposable* (ccND) if for all *x*^1^, *x*^2^ ∈ *P* with sign(*x*^1^), sign(*x*^2^) ≤ sign(*x*) and 0 < λ < 1, the decomposition *x* = λ*x*^1^ + (1 – λ)*x*^2^ implies *x*^1^ = *x*^2^.

Let 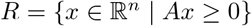 be the *recession cone* of *P*. A vector *e* ∈ *P* ∪ *R* is *elementary* (an EV of *P*) if *e* ∈ *P* is ccND or *e* ∈ *R* is cND.

Ultimately, Theorem 10 can be extended to general polyhedral cones. See [11, Theorem 13].

##### Theorem 12.

*Let P* = { *x* | *Ax* ≥ *b*} *be a polyhedron and R* = {*x* | *Ax* ≥ 0} *its recession cone. Every vector x* ∈ *P is a conformal sum of EVs. That is, there exist finite sets E*_0_ ⊆ *R and E*_1_ ⊆ *P of EVs such that*

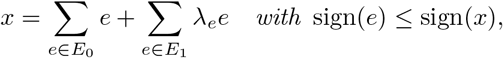

λ_*e*_ ≥ 0, *and* 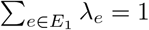. (*Hence*, |*E*_1_| ≥ 1.)

*The set E* = *E*_0_ ∪ *E*_1_ *can be chosen such that* |*E*| ≤ dim(*P*) + 1 *and* |*E*| ≤ | supp(*x*)| + | supp(*Ax*)| + 1.

### 3.2 Software

Simulations were performed in `Python` 3.7.9 using the package `efmtool` 0.2.0 [27]. Fig 1 was created with `BioRender.com`. The remaining figures were created with `R` 4.0.2. All code is available on `GitHub` at https://github.com/diana-sz/EGMs.

## 4 Discussion

In traditional models of cellular growth, elementary flux modes and vectors (EFMs [4, 5] and EFVs [6, 7]) allow the unbiased characterization of all feasible metabolic phenotypes in terms of unique functional units, namely EFMs (in flux cones) and EFVs (in flux polyhedra). More specifically, a certain cellular function can only be performed if there is a corresponding EFM or EFV. However, EFMs and EFVs depend on a fixed biomass composition, which is specified as a biomass “reaction” in addition to the actual metabolic reactions. As a consequence, EFMs and EFVs are not able to describe cellular resource reallocations, in particular, upon changes of environmental conditions [45–47].

In more comprehensive models, the biomass reaction is replaced by explicit synthesis reactions for all (model-specific) macromolecules. In such next-generation models (as used in RBA [3]), biomass composition is no longer fixed, but results from the expression of all proteins (and the synthesis of other macromolecules). In fact, the concentrations *x* = *Nv/μ* of all molecular species are determined by the stoichiometric matrix *N*, the fluxes *v*, and growth rate *μ*, cf. Eq (4).

In this work, we developed a mathematical theory that allows the unbiased characterization not only of all feasible metabolic phenotypes, but also of all feasible biomass compositions in comprehensive, next-generation models. In analogy to EFMs and EFVs, we introduced elementary growth modes and vectors (EGMs and EGVs) which can be interpreted as unique functional units of self-fabrication.

Since EGMs are the basic “building blocks” of any possible flux distribution in *purely stoichiometric* models of cellular growth, they are typically not “self-fabricating” themselves. In particular, they do not reflect implications from autocatalysis, namely that all catalysts of active reactions are synthesized. Still, additional capacity constraints of the form *v_E_* ≤ *k*_cat_ *x_E_* ensure nonzero enzyme concentrations *x_E_* for nonzero fluxes *v_E_*. As a consequence, EGVs in *constraint-based* models are often autocatalytic, as in our running example.

### Comparison with traditional models

It is elucidating to explicitly compare our theory for comprehensive growth models with the analysis of traditional models. In the running example, we take the metabolic part of the stoichiometric matrix (shaded in blue in Fig 1) and add a fixed biomass composition as the last column; see Fig 8 for the resulting stoichiometric matrix. In fact, we choose the biomass composition at maximum growth rate (computed for the comprehensive growth model), cf. Fig 4. Instead of the concentrations of the individual enzymes and the ribosome (which are not part of the traditional model), we use their amino-acid content. The total amino acid concentration is given by

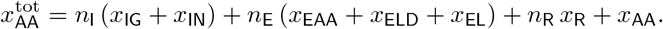

**Fig 8.**
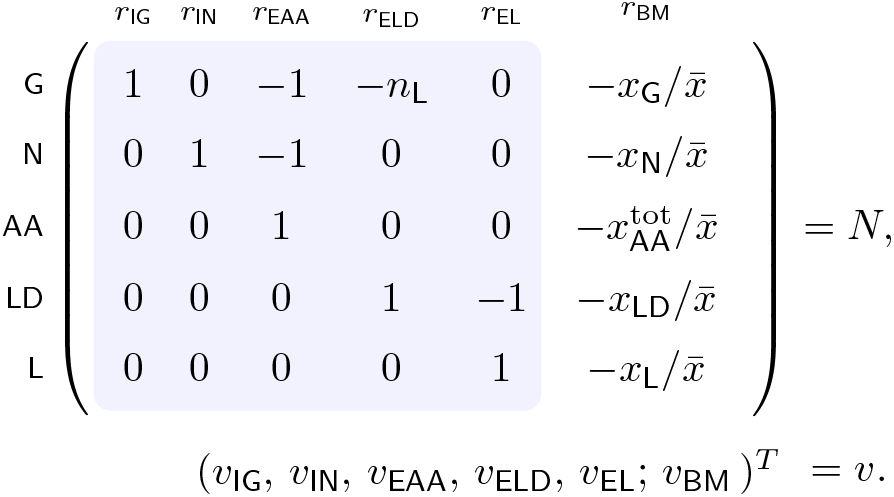
The stoichiometric matrix and the flux vector for a traditional model of the running example. The matrix consists of the metabolic part (shaded in blue, cf. Fig 1) and the biomass composition (last column). Scaling with the concentration 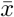 is required to obtain dimensionless entries.

Additionally, we consider enzyme capacity constraints as given in (7a) using the enzyme concentrations at maximum growth. We obtain one EFM and one EFV in the constraint-based model. In other words, the traditional model is feasible, but does not provide further information. This is in stark contrast to the comprehensive model, where we identified 11 EGMs and 10 (or 24) EGVs in the constraint-based model (depending on growth rate), cf. Subsections 2.3 and 2.6. Most importantly, our EGM/EGV analysis provided a characterization of all feasible biomass compositions. In biological terms, we showed that the experimentally observed carbon accumulation upon nitrogen starvation is predicted by our small model of a self-fabricating cell.

### Comparison with (semi-)kinetic models

Recently, de Groot *et al*. [9] analyzed *(semi-)kinetic* models of cellular growth, and one of the authors of this work started a study of constraint-based and (semi-)kinetic models; for a preprint, see [48]. Here, we compare the theoretical concepts introduced in [9] and in this work by means of an example, namely, the minimal growth model shown in Fig 9(a). The following discussion is based on a complete mathematical analysis given in Supporting Information D.

**Fig 9.**
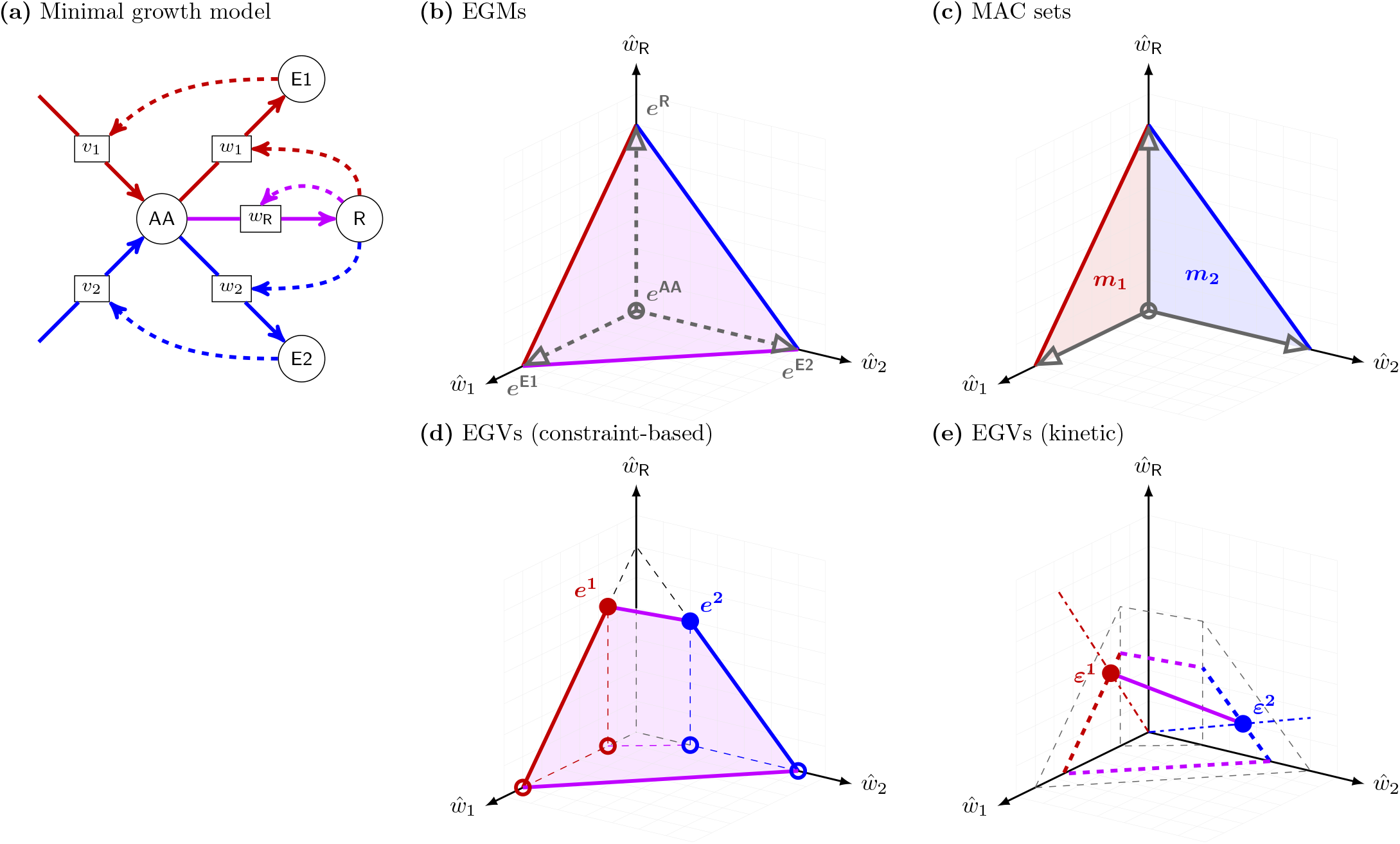
Comparison of EGMs, MAC sets, and EGVs. **(a)** The minimal growth model with alternative pathways is given by 4 molecular species {AA, E1, E2, R} (circles) and 5 reactions (rectangles) labeled by the corresponding fluxes *v* = (*v*_1_, *v*_2_; *w*_1_, *w*_2_, *w*_R_)^*T*^. The “enzymes” E1, E2 catalyze the formation of amino acids AA, and the ribosome R catalyzes the synthesis of the enzymes and the ribosome itself. **(b)** After scaling 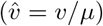 and projection to the (scaled) synthesis fluxes 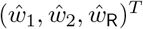, the growth cone becomes a growth polytope (a 3-dimensional simplex), generated by the (scaled and projected) EGMs *e*^AA^, *e*^E1^, *e*^E2^, and *e*^R^. Every EGM produces exactly one species as indicated by its name. **(c)** After projection, the MAC subsets of reactions are *m*_1_ = {*w*_1_, *w*_R_} and *m*_2_ = {*w*_2_, *w*_R_} (indicated by the corresponding fluxes). MAC sets represent minimal pathways: GMs with support *m*_1_ (in red) lie in the two-dimensional simplex generated by the EGMs *e*^AA^, *e*^E1^, and *e*^R^. Analogously for GMs with support *m*_2_ (in blue). EGMs (in gray), lying on the axes, are not AC. **(d)** Enzyme capacity imposes additional *inequality* constraints, and the growth polytope gets restricted (depending on growth rate and *k*^cat^ values). Here, the growth polytope is generated by 6 (scaled and projected) EGVs. Thereby, only EGVs *e*^1^ and *e*^2^ have nonzero ribosome flux and hence are AC. **(e)** Alternatively, in (semi-)kinetic models, enzyme kinetics imposes additional *equality* constraints, and the growth polytope gets restricted (depending on growth rate and metabolite concentrations). Here, the growth polytope is generated by the (scaled and projected) EGVs *ε*^1^ and *ε*^2^ (and hence is 1-dimensional). The EGVs have nonzero ribosome flux and hence are AC. For a complete mathematical analysis of the minimal growth model, see Supporting Information D.

In the example, a cell is able to grow on alternative substrates. Both “enzymes” E1 and E2 can form the amino acids AA used by the ribosome R to synthesize the enzymes and the ribosome itself. Recall that the growth cone is the set of all stoichiometrically feasible fluxes that allow growth (*Nv* ≥ 0 instead of *Nv* = 0 in the classical setting). After scaling and projection to the synthesis fluxes (*w*_1_, *w*_2_, *w*_R_)^*T*^, the growth cone becomes a growth polytope (a 3-dimensional simplex) generated by the (scaled and projected) EGMs *e*^AA^, *e*^E1^, *e*^E2^, and *e*^R^, see Fig. 9(b). This is in analogy to the classical setting, where EFMs generate the flux cone.

However, EGMs are not autocatalytic (AC). In the example, each EGM produces exactly one molecular species, but not all catalysts of active reactions are synthesized. For instance, the EGM *e*^E1^ produces E1 (in reaction *w*_1_), but the ribosome R that catalyzes *w*_1_ is not produced. In technical terms, *e*^E1^ is basically catalytic (BC), since it contains an active catalytic reaction, but not catalytically closed (CC). Only if we combine *e*^E1^ with *e*^R^ (producing the ribosome in reaction *w*_R_), the resulting flux becomes CC and hence AC. In fact, every (positive) combination of *e*^E1^, *e*^R^ and *e*^AA^ is AC, and its support *m*_1_ = {*w*_1_, *w*_R_} is a minimally autocatalytic (MAC) set of reactions. Such fluxes represent minimal pathways and lie on a 2-dimensional facet of the growth polytope, see Fig 9(c).

Additional enzyme capacity constraints restrict the growth polyhedron. In the example, the growth polytope is generated by 6 (scaled and projected) EGVs, see Fig 9(d). All EGVs fulfill the enzyme capacity constraints and hence are CC w.r.t. the enzymatic reactions. However, only EGVs *e*^1^ and *e*^1^ have nonzero ribosome flux and hence are fully AC. Still, every biologically meaningful flux (fulfilling the constraints) is a (nonnegative) combination of EGVs. Again, this is in analogy to the classical setting, where EFVs generate the flux polyhedron [7], determined by a fixed biomass composition, cf. Fig 8. However, in comprehensive models such as in this example, every element of the growth polytope corresponds to a different biomass composition, which allows to study the re-allocation of cellular resources.

In (semi-)kinetic models, enzyme kinetics (for fixed metabolite concentrations) further restricts the growth polytope. In the example, the EGVs *ε*^1^ and *ε*^2^ generate the growth polytope (a 1-dimensional simplex), see Fig 9(e). Given that enzyme kinetics and enzyme capacity constraints use the same *k*^cat^ values, the growth polytope for the kinetic model lies inside the polytope for the constraint-based model. Both EGVs have nonzero ribosome flux and hence are AC.

In the light of this discussion, we find that “elementary growth states” as introduced in de Groot *et al*. [9] are (projections of) EGVs in (semi-)kinetic models. Equivalence classes of their supports (called “elementary growth modes” in [9]) are MAC sets of reactions. In *our* definition, EGMs are the elementary vectors of the growth cone (in the sense of polyhedral geometry). They only depend on stoichiometry (but not on kinetics) and are defined for arbitrary models (including detailed synthesis reactions for macromolecules).

### Elementary vectors and computation

Whereas EFMs are *support-minimal* vectors of the flux cone (a special cone), already EFVs are *conformally non-decomposable* (and not support-minimal) vectors of a flux polyhedron. Also EGMs of the growth cone (a general polyhedral cone) and EGVs of a growth polyhedron are conformally non-decomposable vectors. Thus, conformal non-decomposability (and not support-minimality) is the key feature that defines elementary vectors in computational models of cellular growth, see also [7].

In terms of computation, EGM and EGV analysis (just like EFM and EFV analysis) suffer from a combinatorial explosion in the number of elementary vectors with the size of the network. As with EFMs and EFVs, one may focus on the enumeration of subsets of EGMs and EGVs or consider extra constraints such as thermodynamic feasibility [49, 50].

## Acknowledgments

We thank Christoph Flamm and Ralf Steuer for discussions on a previous version of the manuscript.

SM was supported by the Austrian Science Fund (FWF), project P33218. DS and JZ acknowledge support from the Austrian Center of Industrial Biotechnology (acib). The COMET center *acib* – *Next Generation Bioproduction* is funded by BMK, BMDW, SFG, Standortagentur Tirol, Government of Lower Austria, and Vienna Business Agency in the framework of COMET Competence Centers for Excellent Technologies. The COMET Funding Program is managed by the Austrian Research Promotion Agency (FFG).

The funders had no role in study design, data collection and analysis, decision to publish, or preparation of the manuscript.

## Supporting Information

### A The dynamic model of cellular growth

We derive the dynamic model of cellular growth (1), studied in the main text.

We denote fundamental objects and quantities as follows:

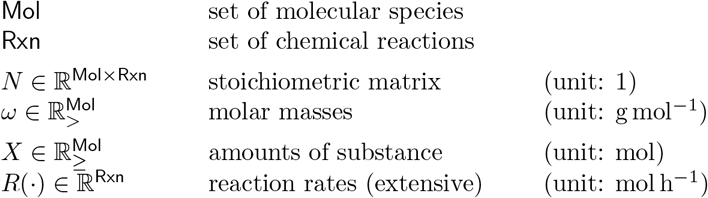

The chemical reactions induce the dynamical system

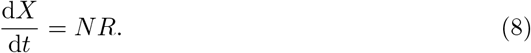

We define mass,

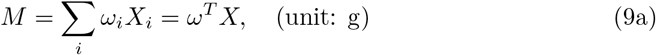

the intensive quantities

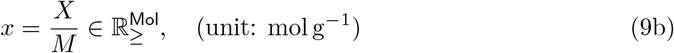

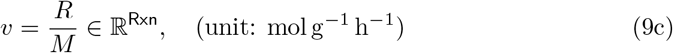

and growth rate

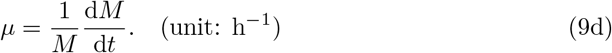

Thereby, we use mass instead of volume to define the “concentrations” *x*, the (intensive) reaction rates *v*, and growth rate *μ*. In practice, cellular composition is often given in the unit molg^−1^ (dry weight).

Finally, we recall the chain rule (of differentiation),

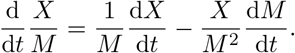

Equations (8), (9), and the chain rule yield the dynamic model of cellular growth:

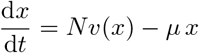

and

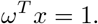

Thereby, we assume constant cell density. Recall that reaction rates depend on (volumetric) concentrations *X/V*,

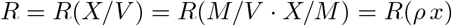

with volume *V* (unit: L) and cell density 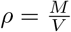 (unit: gL^−1^). Hence,

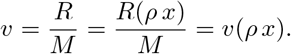

For constant cell density *ρ*, *v* = *v*(*x*) only depends on concentrations.

For alternative derivations, see e.g. [2] or [1].

By multiplying the mass balance equation with a vector 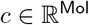, we obtain

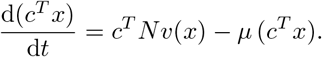

We highlight two observations that hold for any model of cellular growth.

#### Fact (conservation laws)

In a model of cellular growth, there cannot be any conservation laws. In mathematical terms, ker 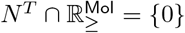.

To see this, assume *c^T^N* = 0 with 0 ≠ *c* ≥ 0, for example, assume *c*_1_ = *c*_2_ = 1 and *c_i_* = 0, otherwise. Then, 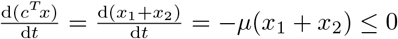, and *μ* > 0 implies *x*_1_ = *x*_2_ = 0 at steady state.

#### Fact (dependent concentrations)

In a model of cellular growth, there can be dependent concentrations. In mathematical terms, ker *N^T^* ≠ {0}.

To see this, assume *c^T^N* = 0 with 0 ≠ *c*, for example, assume *c*_1_ = 1, *c*_2_ = –1, and *c_i_* = 0, otherwise. Then, 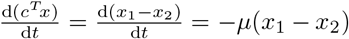, and *μ* > 0 implies *x*_1_ = *x*_2_ at steady state.

### B Example: membrane constraints

For the small model of a self-fabricating cell studied in the main text, we derive the membrane constraints (7c) and (7d).

The cell membrane area *A* is formed by lipids L and importers IG and IN,

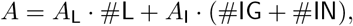

where *A*_L_ and *A*_I_ denote the areas of lipids and importers, respectively, and #*X* denotes the number of molecule *X*. After division by Avogadro’s number *N_A_*, we have

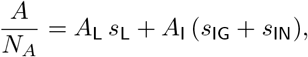

where 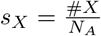 denotes the amount of substance. Further, after division by cell mass *m*, we have

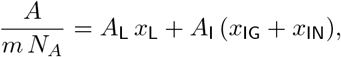

where 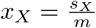 denotes the (mass-specific) concentration. Finally, using cell volume *V*, the surface-to-volume ratio 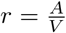, and cell density 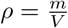, we obtain 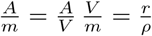 and hence

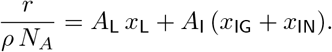

Additionally, we require that a minimum fraction *α* of the surface area is formed by lipids,

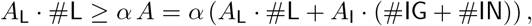

that is,

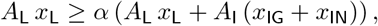

where we use concentrations instead of numbers of molecules. Equivalently,

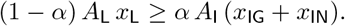

### C Example: figures and tables

**Table S1.**
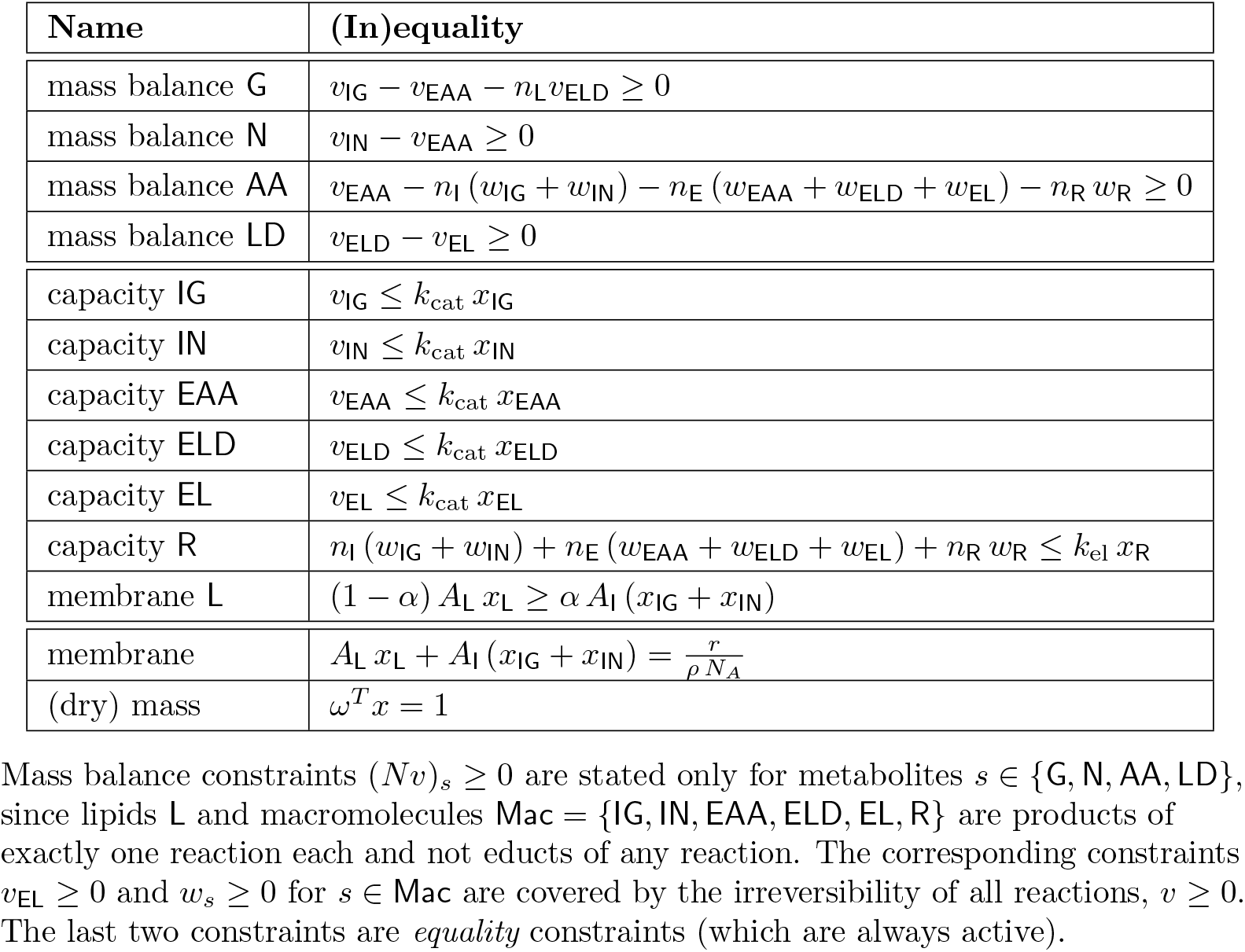
Essential constraints for the small model of a self-fabricating cell.

| Name | (In)equality |
| --- | --- |
| mass balance G | $v_{\text{IG}} - v_{\text{EAA}} - n_{\text{L}} v_{\text{ELD}} \geq 0$ |
| mass balance N | $v_{\text{IN}} - v_{\text{EAA}} \geq 0$ |
| mass balance AA | $v_{\text{EAA}} - n_{\text{I}} (w_{\text{IG}} + w_{\text{IN}}) - n_{\text{E}} (w_{\text{EAA}} + w_{\text{ELD}} + w_{\text{EL}}) - n_{\text{R}} w_{\text{R}} \geq 0$ |
| mass balance LD | $v_{\text{ELD}} - v_{\text{EL}} \geq 0$ |
| capacity IG | $v_{\text{IG}} \leq k_{\text{cat}} x_{\text{IG}}$ |
| capacity IN | $v_{\text{IN}} \leq k_{\text{cat}} x_{\text{IN}}$ |
| capacity EAA | $v_{\text{EAA}} \leq k_{\text{cat}} x_{\text{EAA}}$ |
| capacity ELD | $v_{\text{ELD}} \leq k_{\text{cat}} x_{\text{ELD}}$ |
| capacity EL | $v_{\text{EL}} \leq k_{\text{cat}} x_{\text{EL}}$ |
| capacity R | $n_{\text{I}} (w_{\text{IG}} + w_{\text{IN}}) + n_{\text{E}} (w_{\text{EAA}} + w_{\text{ELD}} + w_{\text{EL}}) + n_{\text{R}} w_{\text{R}} \leq k_{\text{el}} x_{\text{R}}$ |
| membrane L | $(1 - \alpha) A_{\text{L}} x_{\text{L}} \geq \alpha A_{\text{I}} (x_{\text{IG}} + x_{\text{IN}})$ |
| membrane | $A_{\text{L}} x_{\text{L}} + A_{\text{I}} (x_{\text{IG}} + x_{\text{IN}}) = \frac{r}{\rho N_{\text{A}}}$ |
| (dry) mass | $\omega^T x = 1$ |

Mass balance constraints $(Nv)_s \geq 0$ are stated only for metabolites $s \in \{\text{G}, \text{N}, \text{AA}, \text{LD}\}$ , since lipids **L** and macromolecules $\text{Mac} = \{\text{IG}, \text{IN}, \text{EAA}, \text{ELD}, \text{EL}, \text{R}\}$ are products of exactly one reaction each and not educts of any reaction. The corresponding constraints $v_{\text{EL}} \geq 0$ and $w_s \geq 0$ for $s \in \text{Mac}$ are covered by the irreversibility of all reactions, $v \geq 0$ . The last two constraints are *equality* constraints (which are always active).

**Fig S1.**
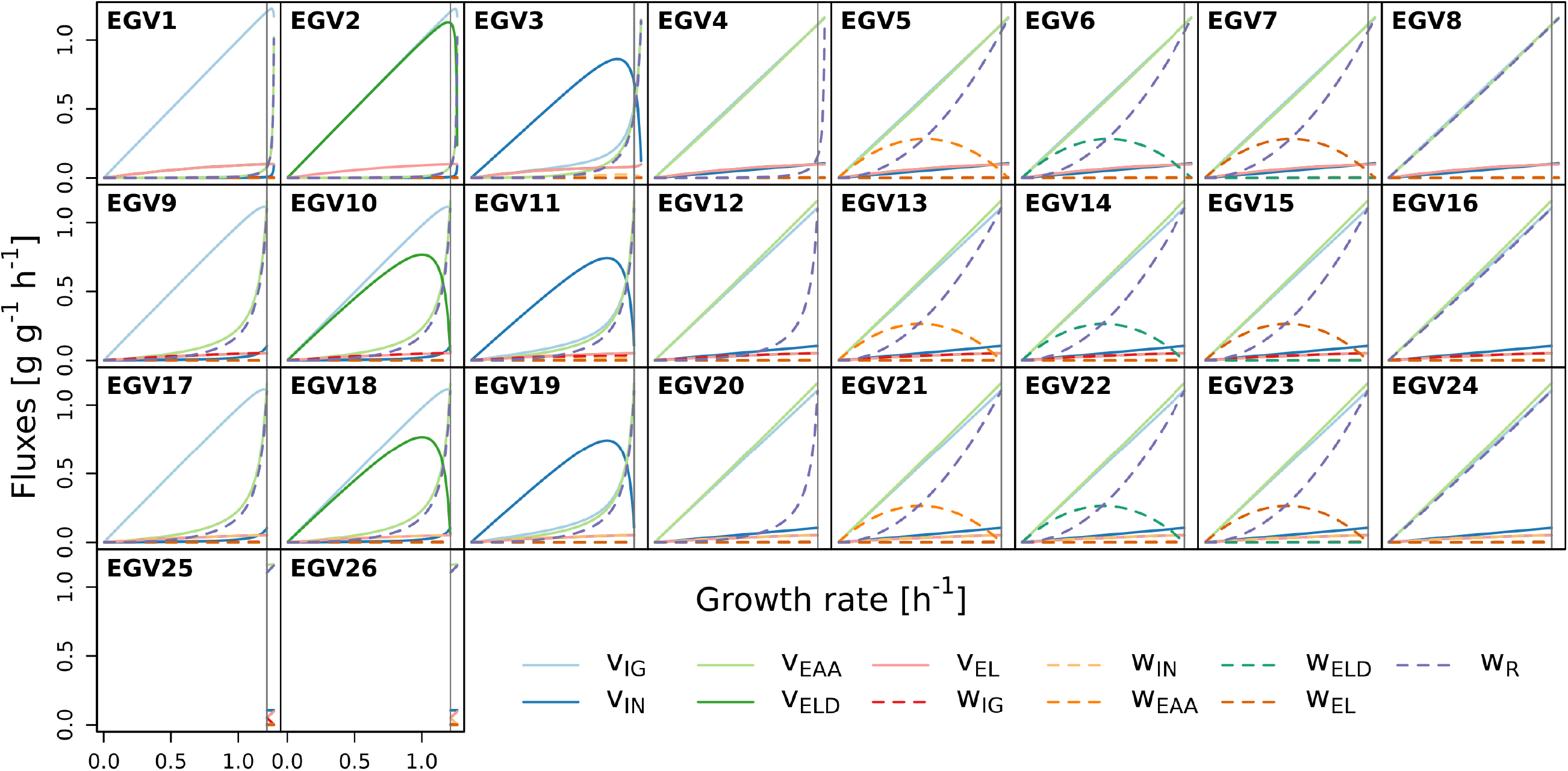
(Scaled) fluxes for all 26 EGVs.

**Fig S2.**
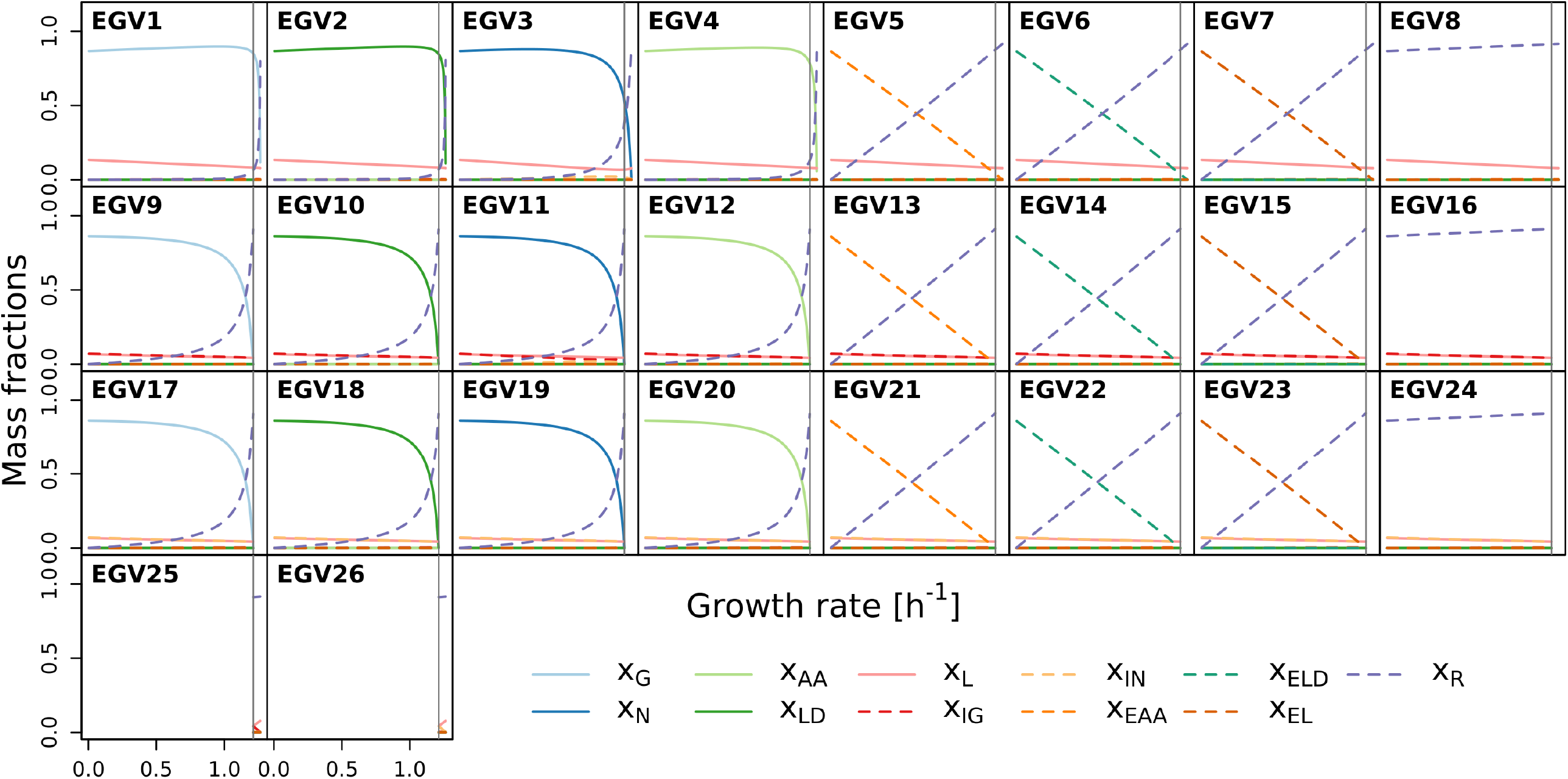
Mass fractions for all 26 EGVs.

**Fig S3.**
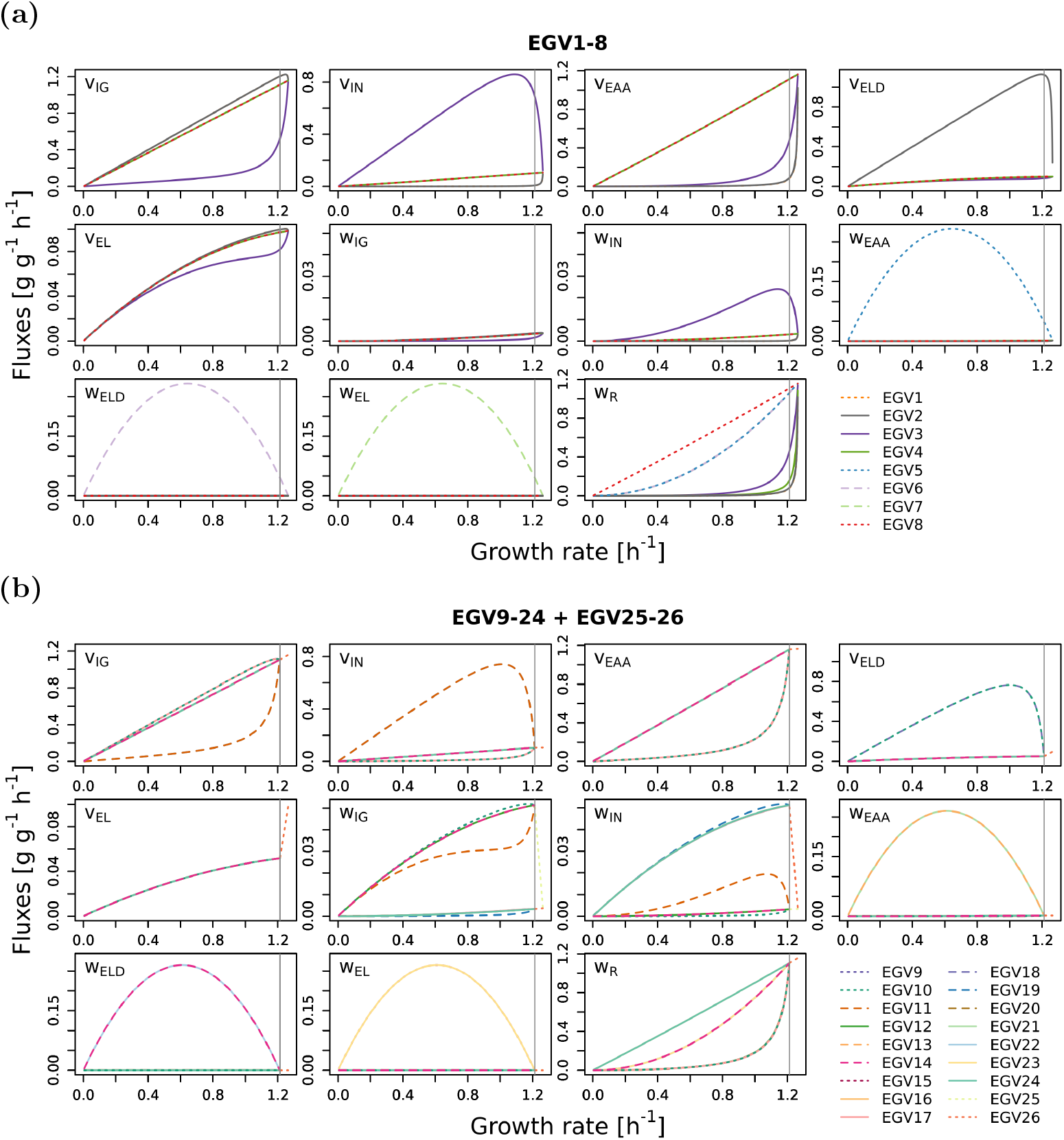
(Scaled) fluxes for (a) the 8 EGVs that exist for all growth rates and (b) the 16 EGVs that exist in regime L plus the 2 EGVs that exist in regime H.

### D Minimal growth model with alternative pathways

Consider the following minimal model of cellular growth with two alternative pathways:

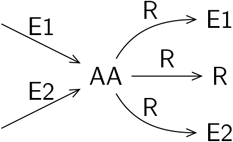

The cell takes up external substrates and forms amino acids (AA) via two “reactions”,

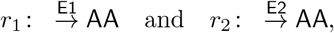

catalyzed by the “enzymes” E1 and E2, respectively. Amino acids are then used by the ribosome (R) to synthesize the enzymes and the ribosome itself,

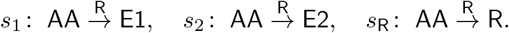

The set of molecular species is Mol = {AA, E1, E2, R}, and the set of reactions is Rxn = Rmet ∪ Rsyn with metabolic reactions Rmet = {*r*_1_, *r*_2_} and synthesis reactions Rsyn = {*s*_1_, *s*_2_, *s*_R_}.

The resulting stoichiometric matrix and the corresponding flux vector are given by

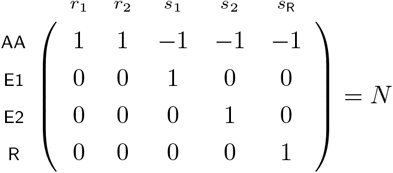

and

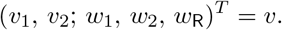

By mass conservation (for the synthesis reactions), the molar masses obey 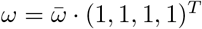. The growth cone is given by

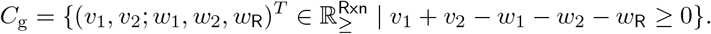

As it turns out, there are 4 × 2 = 8 EGMs (up to scaling), corresponding to the four species and the two alternative pathways, that is, 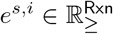 with *s* ∈ Mol and *i* ∈ {1, 2}. Explicitly,

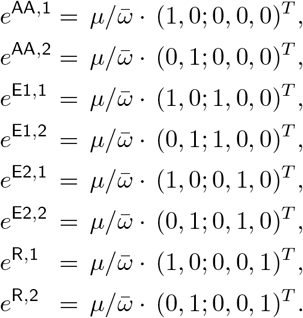

Every EGM “produces” exactly one molecular species, as indicated by its name, thereby using either pathway 1 or 2. (For every EGM, there is exactly one species with nonzero associated concentration.) Due to the factor 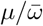, all EGMs have associated growth rate *μ*.

GMs need not be AC, in particular, no EGM is AC. In fact, every (nonzero) GM is BC, since all reactions are catalytic, however, a GM need not be CC. MAC subsets of reactions are the supports of AC GMs. There are two MAC sets, namely *M*_1_ = {*r*_1_, *s*_1_, *s*_R_} and *M*_2_ = {*r*_2_, *s*_2_, *s*_R_}, corresponding to the two alternative pathways. AC GMs with support *M*_1_ are generated by the EGMs *e*^AA,1^, *e*^E1,1^, *e*^R,1^. (Analogously for the MAC set *M*_2_.)

For a GM *v* ∈ *C*_g_, the associated growth rate amounts to

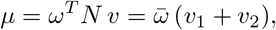

determined by the “exchange” fluxes *v*_1_ and *v*_2_. For fixed growth rate *μ*, the growth cone becomes a growth polytope,

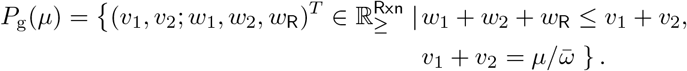

Further, for scaled fluxes 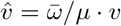, the polytope becomes independent of *μ*,

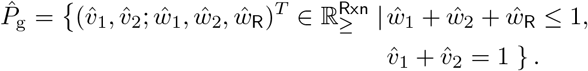

In particular, its projection to the synthesis fluxes 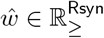 is the “growth simplex”

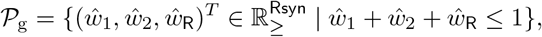

spanned by the projections of the scaled EGMs,

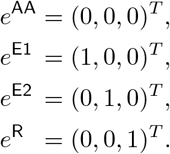

The “projections” of the MAC sets *M*_1_ and *M*_2_ to the synthesis reactions are *m*_1_ = {*s*_1_, *s*_R_} and *m*_2_ = {*s*_2_, *s*_R_}. Scaled projections of AC GMs with support *m*_1_ lie in the (two-dimensional) simplex generated by *e*^AA^, *e*^E1^, *e*^R^. (Analogously for *m*_2_.)

Catalytic closure can be ensured by additional constraints.

- In a constraint-based model, one considers *inequality* enzyme capacity constraints,

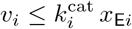

for *i* ∈ {1, 2}, whereas
- in a (semi-)kinetic model, one considers *equality* constraints arising from enzyme and ribosome kinetics,

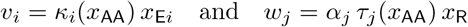

for *i* ∈ {1, 2} and *j* ∈ {1, 2, R}. Thereby, *κ_i_*, *τ_j_* are functions of the amino acid concentration *x*_AA_, and *α_j_* are control parameters (ribosome fractions) for studying growth rate maximization, cf. [1, 3].

Moreover, one often considers ribosome capacity constraints: *w*_1_ + *w*_2_ + *w*_R_ ≤ *k*_tl_ *x*_R_ in constraint-based models and 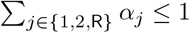 in (semi-)kinetic models. However, the (inequality) ribosome capacity constraint is treated separately in the (semi-)kinetic model, and for reasons of comparison, we just require *x*_R_ > 0 in the constraint-based model.

Additional constraints involve concentrations. For a GM *v* ∈ *C*_g_, the associated concentrations *x* are given by

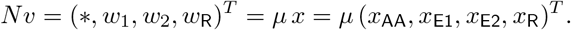

In particular, *x*_E1_ = *w*_1_/*μ* and *x*_E2_ = *w*_2_/*μ*.

In the following, we consider the growth polyhedra and EGVs arising from the additional constraints.

#### Constraint-based model

For fixed growth rate *μ* > 0, the growth polytope *P*_g_ (*μ*) above is further restricted by *inequality* constraints,

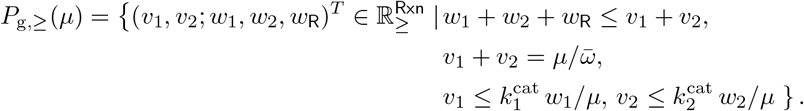

For scaled fluxes 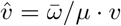, the polytope becomes

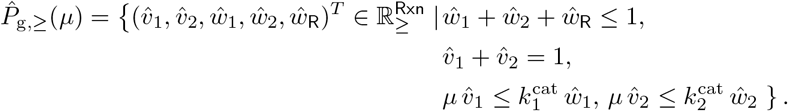

Its projection to the synthesis fluxes 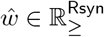 is given by

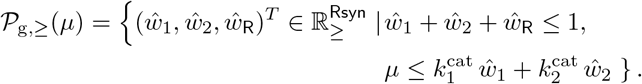

(Scaled) EGVs are the ccND vectors of a (scaled) growth polytope. (Here, since 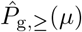 is contained in the nonnegative orthant, EGVs are the vertices.) The number of EGVs depends on *μ*. For small *μ*, there are 8 EGVs. As it turns out, there are at most two EGVs with nonzero ribosome concentration/flux, namely

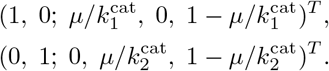

Only these EGVs can fulfill the additional ribosome capacity constraint. They are AC, and their supports are the MAC sets *M*_1_ and *M*_2_. Their projections are

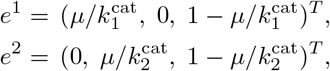

which have supports *m*_1_ and *m*_2_, respectively.

#### (Semi-)kinetic model

For fixed growth rate *μ* > 0 and fixed amino acid concentration 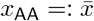 (and hence fixed functions 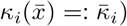, the growth polytope *P*_g_(*μ*) is further restricted by *equality* constraints,

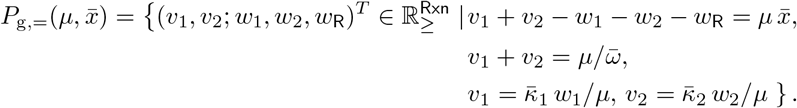

For scaled fluxes 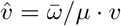, the polytope becomes

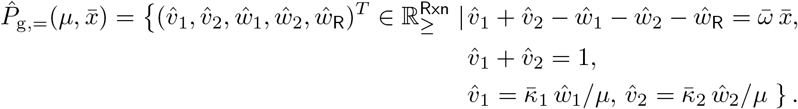

Since 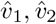 depend on 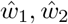, the polytope is in one-to-one correspondence with its projection to the synthesis fluxes 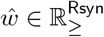,

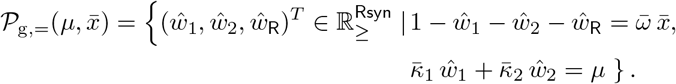

Again, scaled EGVs are vertices of the scaled polytope 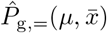. As it turns out, there are at most two EGVs, in particular, at most two EGVs with nonzero ribosome concentration/flux, namely

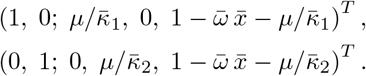

These EGVs are AC, and their supports are the MAC sets *M*_1_ and *M*_2_. They are in one-to-one correspondence with their projections,

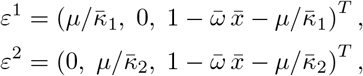

the vertices of the polytope 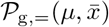, which are called *elementary growth states* (EGSs) in [1]. (In fact, EGSs are defined via the control parameters *α*, which are, however, in one-to-one correspondence with the synthesis fluxes *w*.)

The supports of the EGSs *ε*_1_ and *ε*_2_, that is, *m*_1_ and *m_2_*, the projections of the MAC sets, are called *elementary growth modes* in [1].

